# TGFB1 Induces Fetal Reprogramming and Enhances Intestinal Regeneration

**DOI:** 10.1101/2023.01.13.523825

**Authors:** Lei Chen, Abigail Dupre, Xia Qiu, Oscar Pellon-Cardenas, Katherine D. Walton, Jianming Wang, Ansu O. Perekatt, Wenwei Hu, Jason R. Spence, Michael P. Verzi

## Abstract

The adult gut epithelium has a remarkable ability to recover from damage. To achieve cellular therapies aimed at restoring and/or replacing defective gastrointestinal tissue, it is important to understand the natural mechanisms of tissue regeneration. We employed a combination of high throughput sequencing approaches, mouse genetic models, and murine and human organoid models, and identified a role for TGFB signaling during intestinal regeneration following injury. At 2 days following irradiation (IR)-induced damage of intestinal crypts, a surge in TGFB1 expression is mediated by monocyte/macrophage cells at the location of damage. Depletion of macrophages or genetic disruption of TGFB-signaling significantly impaired the regenerative response following irradiation. Murine intestinal regeneration is also characterized by a process where a fetal transcriptional signature is induced during repair. In organoid culture, TGFB1-treatment was necessary and sufficient to induce a transcriptomic shift to the fetal-like/regenerative state. The regenerative response was enhanced by the function of mesenchymal cells, which are also primed for regeneration by TGFB1. Mechanistically, integration of ATAC-seq, scRNA-seq, and ChIP-seq suggest that a regenerative YAP-SOX9 transcriptional circuit is activated in epithelium exposed to TGFB1. Finally, pre-treatment with TGFB1 enhanced the ability of primary epithelial cultures to engraft into damaged murine colon, suggesting promise for the application of the TGFB-induced regenerative circuit in cellular therapy.

**GRAPHIC ABSTRACT:** 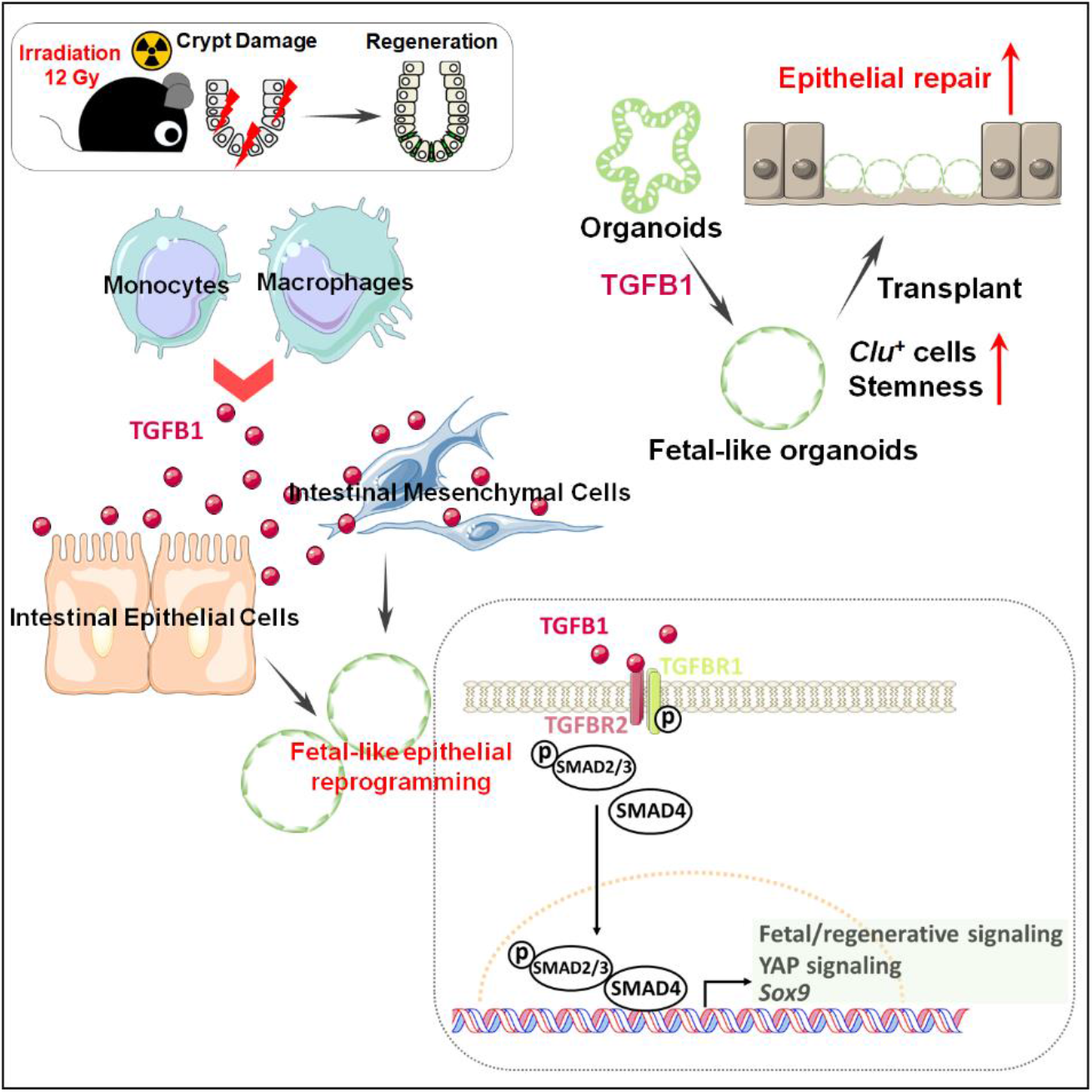

## INTRODUCTION

The intestinal epithelium is a single layer of cells organized into crypts and villi. Under homeostatic conditions, the intestinal epithelium displays a remarkably high turnover rate, where the majority of epithelial cells are replaced every 3 to 5 days, fueled by proliferation of stem and progenitor cells localized in the crypts. The high-rate of crypt cell proliferation makes crypts sensitive to damage by acute inflammation, chemotherapy, or radiotherapy. Upon damage, stem cells are often lost, and differentiated epithelial cell lineages exhibiting plasticity can respond to tissue damage and restore resident intestinal stem cells (ISCs), including crypt base columnar stem cells (CBCs), which are important for regeneration following injury (Metcalfe et al., 2014). For example lineage tracing studies have shown that, *Dll1*^+^ (van Es et al., 2012) and *Atoh1*^+^ secretory (Tomic et al., 2018) progenitor cells, *Lyz1*^+^ Paneth cells (Yu et al., 2018), and *Alpi*^+^ enterocyte progenitor cells (Tetteh et al., 2016) can all de-differentiate to *Lgr5*^+^ ISCs in response to injury, as can cells believed to be slower cycling reserve stem cell populations (Ayyaz et al., 2019; Bala et al., 2022; Okamoto et al., 2020; Roche et al., 2015; Yousefi et al., 2016).

A remarkable property of regenerating intestinal epithelium is its acquisition of a fetal-like transcriptome. The fetal-like, regenerative state is observed in response to helminth infection, irradiation, ablation of CBCs, or DSS-induced colonic damage (Nusse et al., 2018; Yui et al., 2018). Regenerating epithelium in response to virtually all types of tissue damage rely upon activation of the hippo-YAP/TAZ signaling pathway (Ayyaz et al., 2019; Gregorieff et al., 2015; Yui et al., 2018). Transcription factors SOX9 and ASCL2 have also been shown to be important in driving the regenerative response (Murata et al., 2020; Roche et al., 2015), and the fetal-like/regenerative signature appears to be coopted by colon cancers (Bala et al., 2022; Han et al., 2020), further highlighting the importance of this phenomenon. Despite our understanding of cellular plasticity in the intestine, the signaling mechanisms leading to the fetal-like regenerative state and subsequent regenerative process are less well understood.

In this study, we aimed to uncover cells driving the regenerative response, as well as the signaling mechanisms underlying the regenerative ability of intestinal tissues in hopes of finding new therapeutic avenues for intestinal regeneration. Using genetic mouse models, epigenetics, bulk and single cell RNA-sequencing and tissue/organoid culture approaches, we explored the interplay among intestinal epithelial cells, mesenchymal cells and immune cells. We found that TGFB signaling is activated in the intestine post-irradiation, and the main source of TGFB1 is monocytes/macrophages. We further explored the relationship between TGFB signaling and the regenerating intestinal epithelium, as well as the ability of intestinal mesenchyme to promote regeneration after TGFB1 exposure. We discovered a critical role for TGFB1 in promoting intestinal regeneration by inducing a fetal-like, regenerative state in the epithelium via activation of pro-regeneration transcription factors YAP and SOX9. Therapeutically, TGFB1-treated organoids support a more robust tissue engraftment into a mouse model of colitis, suggesting that activation of the regenerative response enhances cellular therapy. These results suggest that modulation of TGFB-signaling could enhance regenerative strategies or cellular therapies in the human intestine.

## RESULTS

### Monocyte/Macrophages deliver TGFB1 ligands to intestinal crypts damaged by irradiation

To better appreciate a time-resolved intestinal response to ionizing radiation, we performed a detailed study across the 5 days post-12 Gy whole body irradiation of mice. Loss of intestinal crypt architecture was widespread by 2 days post-irradiation, with notable loss of proliferative (Ki67 and BrdU incorporation) and stem cell (OLFM4) markers (**Figure 1A-B, Figure S1A**). Between days 2 and 3 post-IR, cellular proliferation in the crypt resumed, most frequently near the crypt-villus junction (**Figure 1B** and **Figure S1A-D**), with highly proliferative, regenerating crypts observed by day 4-5 (**Figure 1A**). Consistent with published reports, transcriptome analysis of isolated crypt epithelium at day 2-3 post-IR identified elevated levels of genes associated with a fetal/regenerative epithelium (Nusse et al., 2018; Wang et al., 2019; Yui et al., 2018); a corresponding decrease was observed in transcripts associated with the crypt-base-columnar stem cell (Munoz et al., 2012) (*Lgr5, Olfm4*, **Figure 1C-D, Figure S1E**). Similar patterns are observed at the single-cell level when comparing crypt epithelial cells from control mice or mice at 3 days post-IR (**Figure 1E**). To specifically focus on cells driving regeneration, we isolated proliferative cells at 56h post-IR, a time point when these cells reappear towards the upper regions of the former crypts. To isolate proliferative cells, we employed *Ki67-RFP* transgenic mice (Basak et al., 2014) and used Fluorescence-Activated Cell Sorting (FACS) followed by scRNA-seq. Compared to Ki67^+^cells in homeostatic crypt epithelium, Ki67^+^ cells within the regenerating epithelium formed a separate cluster when visualized using Uniform Manifold Approximation and Projection (UMAP) and expressed elevated transcript levels of markers associated with fetal/regenerative intestinal epithelium, including *Ly6a* and *Clu* (**Figure 1F-G, Figure S1F**). Taken together, these data indicate that a regenerative response following 12 Gy irradiation initiates around day 2 post-IR with the re-emergence of Ki67^+^ and BrdU^+^ cell populations that express hallmark transcripts of the fetal/regenerative epithelium.

**Figure 1.**
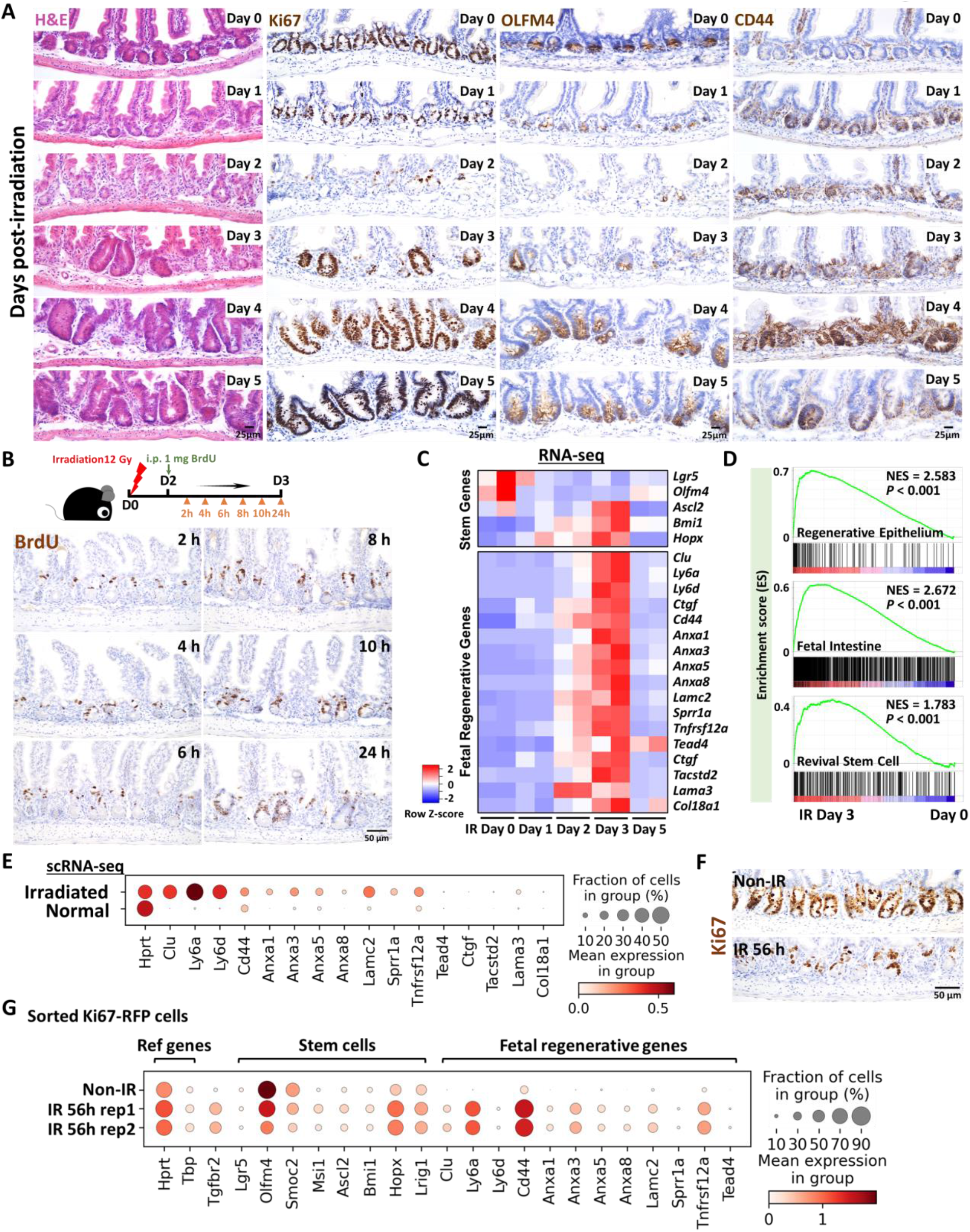
Crypt regeneration mainly starts 3 days after irradiation. **(A)** Demonstration of intestinal regeneration following 12 Gy of irradiation of mice. Crypt cells are lost by 2 days post-irradiation (12 Gy) but restoration begins at Day 3 after irradiation, when highly proliferative regenerative clusters of cells expand, as evidenced by H&E staining and immunohistochemistry staining of stem/proliferative markers (brown color) including Ki67, OLFM4 and CD44 (representative of 3 biological replicates). **(B)** Immunostaining of BrdU (proliferative marker, brown color; representative of 3 biological replicates).Mice were injected with 1 mg of BrdU at Day 2 post-irradiation. Intestinal tissues were harvested after 2, 4, 6, 8, 10 and 24 hours of BrdU injection. **(C)** Heatmap depicts transcript levels of fetal/regenerative marker genes and regenerative stem cell-associated genes that are highly expressed at Day 3 post-irradiation (GSE165157 (Qu et al., 2021), RNA-seq, n=2 biological replicates per time-point). **(D)** GSEA confirms that gene signatures of regenerative epithelium, fetal spheroid and revival stem cells (Ayyaz et al., 2019; Wang et al., 2019; Yui et al., 2018) are elevated at Day 3 post-irradiation (GSE165157 (Qu et al., 2021), crypt cells, n=2 biological replicates per time-course, Kolmogorov-Smirnov test, *P* < 0.001). See expanded panel in **Figure S1E. (E)** scRNA-seq reveals that fetal/regenerative transcripts are elevated in irradiated crypt cells after 3 days of irradiation. Cell numbers per condition (GSE117783 (Ayyaz et al., 2019)) for irradiated crypt cells: n=4252 and normal crypt cells: n=4328. **(F)** Immunostaining of Ki67 after 56 hours of irradiation vs. non-irradiation (proliferative marker, brown color; representative of 3 biological replicates). **(G)** scRNA-seq reveals that fetal/regenerative and reserve stem cell transcripts are elevated in sorted Ki67-RFP positive cells after 56 hours of irradiation. Number of cells in each condition was Non-IR Ki67-RFP positive cells: n=1739; IR 56h Ki67-RFP positive cells replicate 1: n=677; IR 56h Ki67-RFP positive cells replicate 2: n= 669. Ki67-RFP positive cells were isolated and sorted from crypt cells of *Mki67tm1*.*1Cle/J* mice after 56 hours of IR vs. non-IR control. IR: irradiation; Non-IR: non-irradiation (normal control). Also see **Figure S1F**.

We next sought to define key cells and signaling pathways that support the regenerative epithelium. We first conducted scRNA-seq analysis from whole intestinal tissues post-IR, and interrogated ligands known to regulate prominent pathways important for epithelial homeostasis, including the TGFB/BMP and WNT pathways. Of the ligands we analyzed for transcript expression associated with regenerative time points, *Tgfb1* stood out with robust expression, particularly at day 3 post-IR (**Figure 2A**). *Tgfb1* transcripts were enriched in multiple intestinal cell clusters when visualized using UMAP (**Figure 2B, Figure S2A**). We next conducted a time-resolved series of experiments to explore potential TGFB1 contribution to intestinal regeneration (**Figure S2B**). Validation qRT-PCR analysis of *Tgfb1* expression in whole intestine confirmed elevated levels at days 2-3 post-IR, and ELISA assays showed corresponding increase in TGFB1 protein levels at day 3 post-IR in whole intestinal lysates (**Figure 2C-E, Figure S2C-F**).

**Figure 2.**
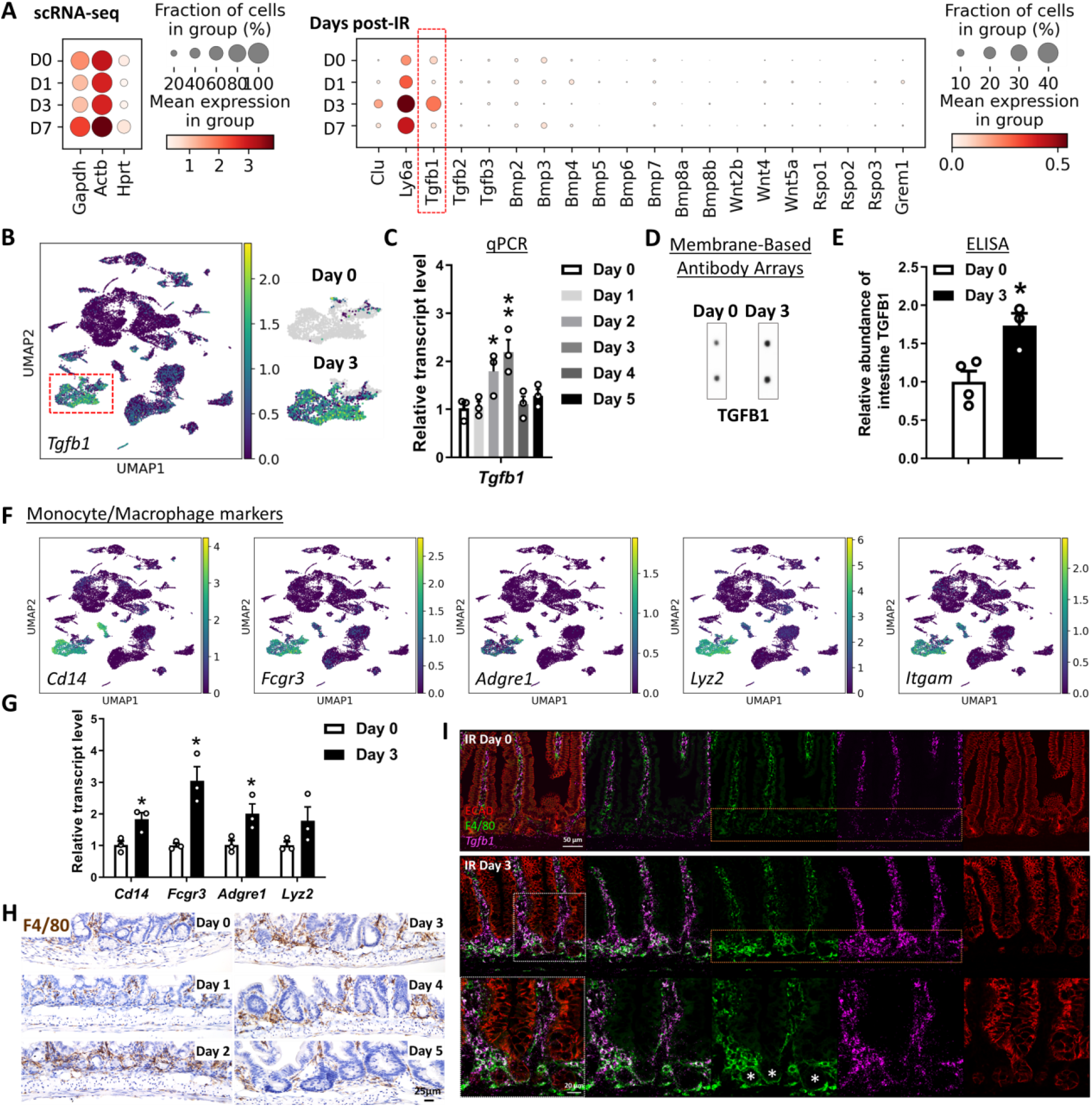
TGFB1 is highly enriched in Day 3 irradiated mouse intestine, and monocytes/macrophages are likely the main cell source of TGFB1. **(A-C)** *Tgfb1* transcript is notably enriched in the intestine at 3 days post-irradiation. **(A)** Dot plots of scRNA-seq data following mouse irradiation at days 0, 1, 3, and 7: GSE165318, duodenum/jejunum boundary, n=3-4 biological replicates per time point) indicate that among secreted regulators of the TGF/BMP/WNT signaling pathways, transcripts corresponding to *Tgfb1* are the most upregulated during regeneration of the gut (red box) and overlap expression of epithelial regenerative markers *Clu* and *Ly6a*. **(B)** UMAP projection of all cells identifies a cell cluster expressing highest levels of *Tgfb1*, particularly at Day 3 post-irradiation. Cells per timepoint: D0: n=4415; D1: n=2995; D3: n=7368; D7: n=3783; D14: n=2220. **(C)** Elevated *Tgfb1* transcript levels are observed at days 2 to 3 post-irradiation. The qRT-PCR data are presented as mean ± SEM (n=3 biological replicates, whole duodenal fragments). Transcript levels are relative to Day 0 (before IR), statistical comparisons were performed using one-way ANOVA followed by Dunnett’s post at *P* < 0.01** or *P* < 0.05*. See schematic of experimental design in **Figure S2B. (D-E)** TGFB1 protein levels are elevated in the intestine at Day 3 post-irradiation. **(D)** Membrane-based antibody arrays: n=2 independent experiments, 2 technical replicates per membrane (See full blots in **Figure S2C**). **(E)** ELISA to measure TGFB1: Data are presented as mean ± SEM (n=3-4 biological replicates, duodenal fragments, Student’s t-test at *P* < 0.05*). **(F)** The same UMAP as in panel B demonstrates that *Tgfb1*-expressing cells co-express markers of monocytes/macrophages. **(G)** qRT-PCR corroborates elevation of transcript levels of monocyte/macrophage marker genes at 3 days post-IR. All qRT-PCR data are presented as mean ± SEM (n=3 biological replicates, duodenal fragments, Student’s t-test at *P* < 0.05*). **(H)** Tissues from mice at different days post-12 Gy irradiation were probed for the monocyte/macrophage marker F4/80 using immunohistochemistry. An increase in F4/80 cells (brown color) occurs when the tissue begins to heal at 2 days post-IR (representative of 3 biological replicates). **(I)** RNAscope localized *Tgfb1* transcripts relative to immunostaining signal from ECAD (epithelial marker) and F4/80 (representative of 3 biological replicates). Co-stains reveal that F4/80-marked macrophages are associated with damaged crypt epithelium at day 3 post-IR and overlap with regions of elevated levels of *Tgfb1*, suggesting that monocyte/macrophages are recruited to the damaged tissue and produce TGFB1.

Increase in TGFB1 corresponded to elevated levels of p-SMAD2/3, the downstream transcriptional effectors of TGFB signaling (**Figure S2G-H**). Further analysis of scRNA-seq data identified a cluster of cells expressing monocyte/macrophage markers as the most prominent source of *Tgfb1* transcripts in the gut post-IR (**Figure 2B**,**F**), and increases in monocyte/macrophage-associated transcripts (**Figure 2G**), protein marker expression (F4/80, **Figure 2H**), and co-expression of macrophage markers and *Tgfb1* transcripts (**Figure 2I**) collectively pointed to monocyte/macrophages being recruited to the damaged crypts and producing TGFB1 ligands at days 2-3 post-IR.

### TGFB1 signaling and monocyte/macrophages are required for epithelial regeneration following irradiation

To further explore a role for TGFB1 in epithelial regeneration, we sought to define TGFB-receptor expression. TGFB receptors are expressed in the epithelium, and *Tgfbr2* is specifically elevated at 3 days post-IR, specifically at a time when homeostatic stem cell markers are reduced and fetal/regenerative marker transcripts are elevated in the epithelium (**Figure 3A, Figure S3A**). *Tgfbr2* transcripts are particularly enriched in epithelial cells co-expressing regenerative markers such as *Clu* and *Ly6a* (**Figure 3B-C, Figure S3A**), and enriched near the crypt zone when assayed *in situ* using RNAscope (**Figure 3D**). Independent scRNA-seq datasets of regenerating intestinal epithelium independently confirm the elevation of *Tgfbr2* in the epithelium following IR and correspond to the pattern of transcripts associated with the regenerative cell state (**Figure 3E, Figure S3B-C**).

**Figure 3.**
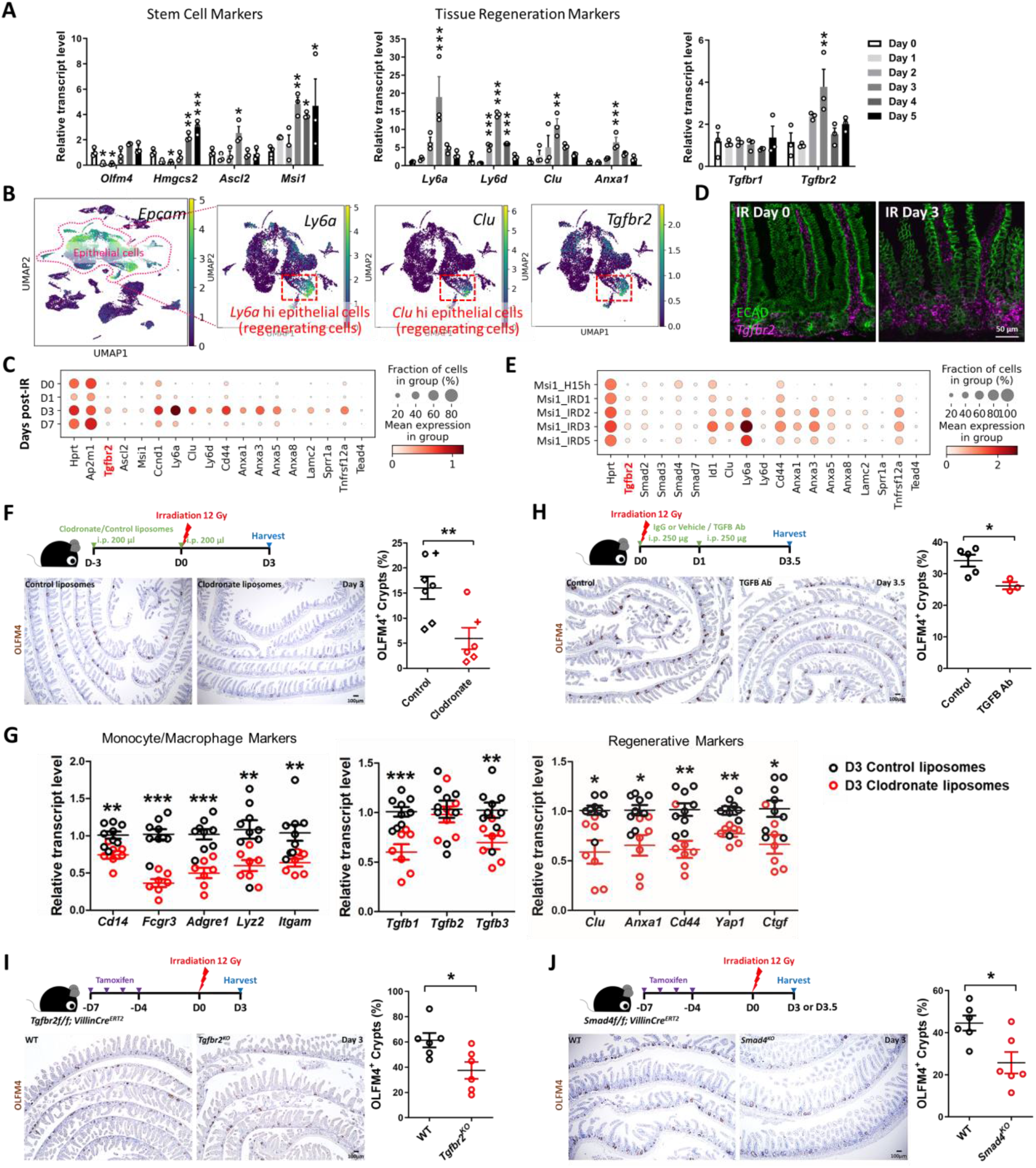
TGFB pathway is required for epithelial regeneration in the intestine after irradiation. **(A)** qRT-PCR analysis indicates that transcript levels of stem cell marker genes, tissue regeneration marker genes and *Tgfbr2* are dynamic in during intestinal regeneration post-irradiation. The qRT-PCR data are presented as mean ± SEM (n=3 biological replicates, duodenal fragments). Transcript levels relative to Day 0 before irradiation. Statistical comparisons were performed using one-way ANOVA followed by Dunnett’s post at *P* < 0.001***, *P* < 0.01** or *P* < 0.05*. **(B)** scRNA-seq of mouse intestines across a time-course post-irradiation (GSE165318). Of all the epithelial cells in the dataset (marked by *Epcam* expression), there is a strong correlation between *Tgfbr2-*expressing cells and the subset of epithelial cells expressing regenerative markers (*Ly6a* and *Clu*, see expanded panel in **Figure S3A**). **(C)** Dot plots of epithelial cells from the same dataset reveal that *Tgfbr2* is highly enriched at Day 3 post-irradiation and correlated with fetal/regenerative gene profiling. *Hprt* and *Ap2m1* were used as reference genes. **(D)** RNAscope reveals elevated *Tgfbr2* transcripts in the Day 3 irradiated intestine (representative of 3 biological replicates). ECAD: epithelial marker. **(E)** An independent scRNA-seq (GSE145866 (Sheng et al., 2020)) dataset also reveals that transcripts related to TGFB pathway and fetal/regenerative genes are elevated at Day 3 post-irradiation in sorted Msi1-GFP positive cells (irradiation-resistant) and their progeny cells. *Msi1-CreERT2; R26-mTmG* mice were treated with tamoxifen for 15 hours and then used as controls or further irradiation for 1, 2, 3, and 5 days. Number of cells in each condition was Msi1_H15h: n= 2281; Msi1_IRD1: n= 1257; Msi1_IRD2: n= 1312; Msi1_IRD3: n= 2989; Msi1_IRD5: n= 1792. **(F)** Monocytes/Macrophages were depleted using clodronate-containing liposomes (2 treatments of 200 μl i.p. injections 72 hours pre- and day of irradiation). Tissues were assessed for regenerative cell clusters using OLFM4 immunostaining. Clodronate-treated samples shows a significant reduction in the number of OLFM4 positive regenerating cell clusters. Different symbols (circle, diamond and cruciform) represent biological replicates from three different batches of experiments (n=6-7 biological replicates, distal duodenum to proximal jejunum, Student’s t-test at *P* < 0.01**). **(G)** qRT-PCR confirms downregulated transcript levels of monocyte/macrophage marker genes, *Tgfb* genes, and regenerative marker genes in the intestine upon clodronate treatment (n=7-9 biological replicates, duodenal fragments, Student’s t-test at *P* < 0.001***, *P* < 0.01** or *P* < 0.05*). Tissues were collected 3 days post-irradiation. **(H)** Mice treated with 2 doses of neutralizing antibodies directed against TGFB were less efficient at regenerating post irradiation compared to control-treated mice, as measured by counting the number of proliferative foci as marked by OLFM4 immunostaining (n=3-5 biological replicates, distal duodenum to proximal jejunum, Student’s t-test at *P* < 0.05*). IgG or vehicle treated mice were used as control mice. Tissues were collected 3.5 days post-irradiation. **(I)** *Tgfbr2* intestine-specific knockout restricts regeneration after irradiation (n=6 biological replicates, duodenum, Student’s t-test at *P* < 0.05*). **(J)** *Smad4* intestine-specific knockout restricts regeneration after irradiation (n=6 biological replicates, Jejunum, Student’s t-test at *P* < 0.05*). Mice were treated with tamoxifen to inactivate *Tgfbr2* or *Smad4* in the intestinal epithelium 7 days before 12 Gy of irradiation. Intestine was collected 3 or 3.5 days post-IR and scored for regenerative foci using OLFM4 immunostaining.

To better appreciate the role of monocyte/macrophages and TGFB1 signaling in intestinal regeneration, we perturbed recruitment of these cells to damaged intestines or blocked TGFB-signaling following 12 Gy irradiation. We depleted monocyte/macrophage populations by treating mice with clodronate-containing liposomes. Clodronate liposome treatment led to a visible reduction in F4/80^+^ cells in intestines 3 days post-IR compared to animals treated with control liposomes (**Figure S3D**), and led to a corresponding decrease in regenerative epithelial foci in the intestine (OLFM4 immunostain, **Figure 3F**). Transcript levels for markers of monocyte/macrophages were also decreased by clodronate treatment, as well as *Tgfb1* transcripts and protein (**Figure 3G, Figure S3E**). A reduction in regenerative cell markers was also observed upon clodronate treatment (**Figure 3G**) consistent with a role for monocyte/macrophage cells in both promoting regeneration and with supplying TGFB1 to the damaged crypts. To specifically target TGFB signaling in the regenerative process, we either treated mice with TGFB-neutralizing antibodies, or genetically inactivated *Tgfbr2* or *Smad4*. In each case, perturbations of the TGFB-signaling pathway led to reduction of regenerative crypts following IR (**Figure 3H-J**). These experiments suggest a mechanism in which monocyte/macrophages promote epithelial regeneration by promoting TGFB1 signaling activity in the intestinal epithelium.

### TGFB1 is necessary and sufficient to induce fetal reversion in intestinal organoid cultures

The previous experiments perform loss-of-function assays to support a role for TGFB1 in driving epithelial regeneration. We next investigated whether exogenous treatment of TGFB1 would be sufficient to induce a regenerative epithelial state in the context of intestinal organoid culture (Sato et al., 2009). We exposed intestinal organoids to 4 Gy IR four days after initiation of crypt culture, and 48 hours later we treated cultures with a single, 24 hour dose of TGFB1 (**Figure 4A**) to approximate the timeline of TGFB1 enrichment in the regenerating gut (**Figures 2-3**). Compared to controls, organoids treated with TGFB1 exhibited a spheroid morphology (**Figure 4B-C**) and a notable elevation in fetal and regenerative cell transcripts (**Figure 4D-E**). TGFB1-induced expression of the fetal/regenerative/revival cell/YAP signaling signatures was i) rapid, occurring within 24 hours, ii) sustained, lasting at least 5 days following the single TGFB1 treatment (**Figure 4F-G, Figure S4A**), iii) dependent upon epithelial *Tgfbr2* expression, as the response did not occur in TGFB1-treated *Tgfbr2*^*KO*^ organoids (**Figure 4G, Figure S4A-C**), and iv) sensitive to TGFBR-inhibitors (**Figure S4D-F**). To better appreciate the timeline of the organoid response to TGFB1 treatment, we performed scRNA-seq analysis of organoids treated with vehicle or TGFB1 for 6, 15, or 24 hours (**Figure 4H**). TGFB1-treatment induced increasingly elevated levels of canonical TGFB1-pathway targets such as *Id1, Smad7*, and *Tgif1* over time, and these increases were accompanied by a marked induction of regenerative markers such as *Clu* and *Anxa*-family genes (**Figure 4I**). Approximately half of the cells expressed transcripts consistent with differentiated enterocytes, and the remainder expressed transcripts associated with progenitor cell populations (**Figure 4J-K**). These analyses point to an induction of the regenerative marker *Clu* as early as 6 hours post-TGFB1 treatment, and maximal expression at the latest time point collected, 24h. RNA velocity analysis and UMAP visualization indicated that *Clu*-expressing cells at the later time points appear most closely related to *Lgr5*-expressing progenitor cells, suggesting a potential origin of these cells. Collectively, these data indicate that TGFB1 potently induces a shift in the morphology and transcriptome of intestinal organoids to acquire properties of regenerating epithelium.

**Figure 4.**
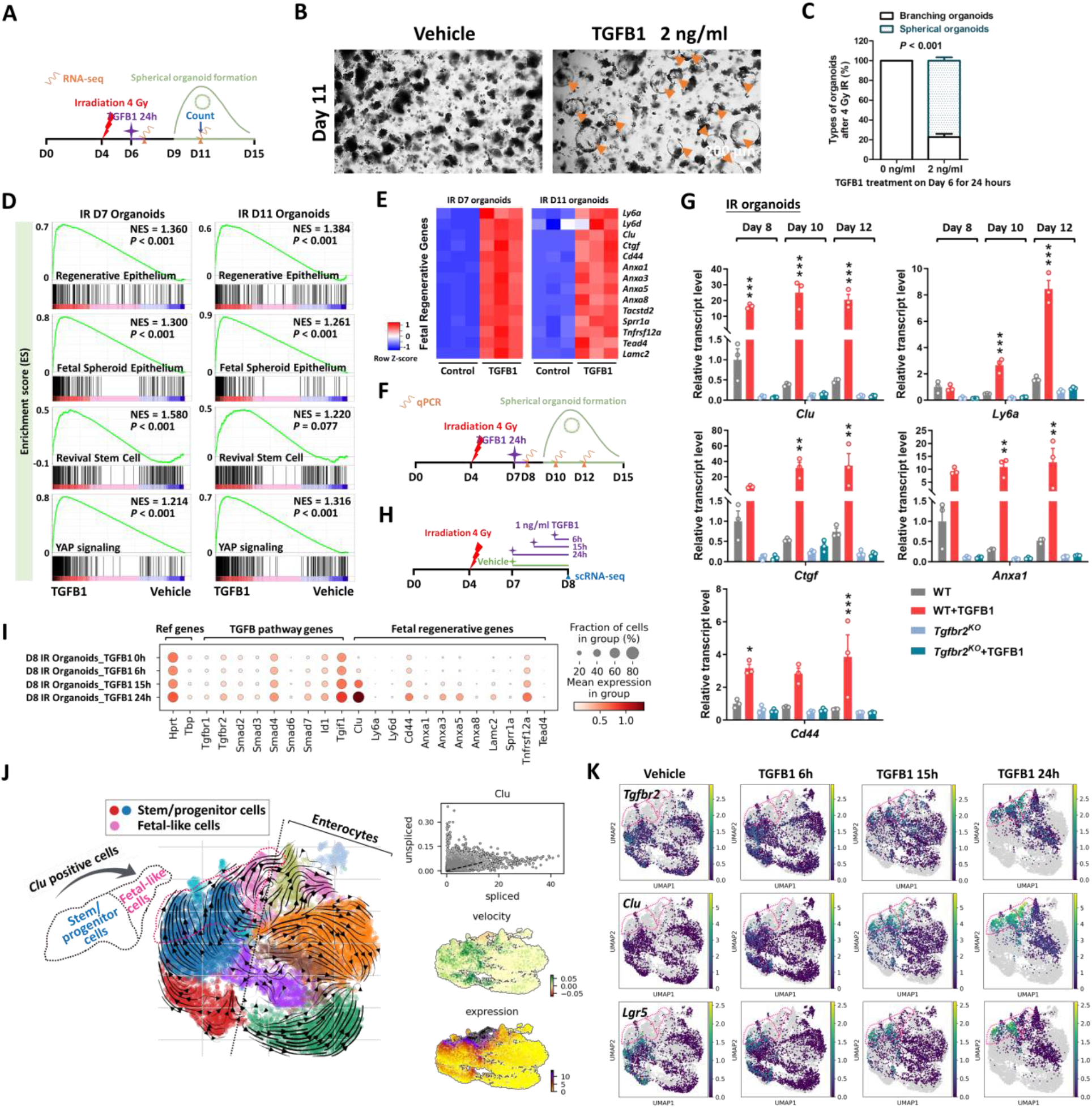
TGFB1 is sufficient to induce fetal/regenerative gene signatures and increase *Clu* positive cells in organoid culture. **(A-E)** Organoids treated with TGFB1 for 24 hours acquire a spheroid morphology (see orange arrows in panel B) and maintain expression of regeneration marker genes for at least 5 days post-TGFB1 treatment. **(A)** Schematic for the experiments to score morphology, counts, and bulk RNA-seq. Primary intestinal organoids were exposed to 4 Gy of irradiation on Day 4, followed by TGFB1 treatment on Day 6 for 24 hours. Organoids were collected for bulk RNA-seq on Day 7 and Day 11. **(B)** Representative images of irradiated organoids treated with vehicle or TGFB1. **(C)** Percentage of branching and spherical organoids upon TGFB1 treatment. Organoids were counted at Day 11, which is 5 days post-TGFB1 treatment (n=3 independent organoid cultures, Student’s t-test at *P* < 0.001). **(D)** Bulk RNA-seq of intestinal organoids cultured with TGFB1 for 24 hours show strong correlation with published gene signatures associated with intestinal regeneration post-DSS injury, fetal spheroid, revival stem cells and YAP signaling (Ayyaz et al., 2019; Gregorieff et al., 2015; Mustata et al., 2013; Wang et al., 2019; Yui et al., 2018), as measured by GSEA (n=3 independent organoid cultures, Kolmogorov-Smirnov test). **(E)** Heatmaps display that RNA-seq expression levels of fetal/regenerative genes are highly expressed upon TGFB1 treatment compared to the vehicle controls (n=3 independent organoid cultures). **(F)** Schematic for the qRT-PCR experimental design. Primary intestinal organoids cultured from *Tgfbr2*^*KO*^ mice (4 days post-tamoxifen) and their littermate WT controls, were exposed to 4 Gy of irradiation on Day 4, followed by TGFB1 treatment (1 ng/ml) on Day 7 for 24 hours. Organoids were collected on Day 8, Day 10 and Day 12 for qRT-PCR. **(G)** qRT-PCR indicates that expression of regeneration marker genes increases within 24 hours and is sustained for at least 5 days post-TGFB1 treatment in the WT organoids, but not in the *Tgfbr2*^*KO*^ organoids. See more examples in **Figure S4A. (H-K)** scRNA-seq of organoids post-irradiation and upon TGFB1 treatment across timepoints. **(H)** Schematic of scRNA-seq experimental design. Primary intestinal organoids were exposed to 4 Gy of irradiation on Day 4, followed by TGFB1 treatment (1 ng/ml) on Day 7. Organoids were collected for scRNA-seq after 6, 15, and 24 hours of TGFB1 treatment. Organoids treated with vehicle were used as control. **(I)** Dot plots show that TGFB1 activates TGFB pathway and fetal/regenerative genes in a time-dependent manner. **(J)** RNA velocity analysis identifies cells in *Lgr5* and *Olfm4*-expressing clusters are synthesizing new *Clu* transcripts. **(K)** UMAP plots indicate that, across time points, there is a close correlation between *Clu* and *Tgfbr2* expression. *Lgr5*-expressing clusters begin to overlap with *Clu*-expressing cells. Number of cells in each condition was Vehicle: n= 2815; TGFB1 6h: n= 4071; TGFB1 15h: n= 2788; TGFB1 24h: n=2177.

### The stromal mesenchyme responds to TGFB1 to promote fetal reversion of the epithelium

scRNA-seq analysis also revealed that *Tgfbr2* transcripts were enriched in *Pdgfra*^+^ mesenchymal cell populations (**Figure 5A, Figure S5A**). *Pdgfra-Lo* cells have demonstrated roles in signaling to the epithelium and regulating growth (Greicius et al., 2018; McCarthy et al., 2020; Shoshkes-Carmel et al., 2018). To define the role of intestinal mesenchyme in promoting TGFB1-dependent regeneration, we cultured *Pdgfra-Lo* mesenchymal fibroblasts from *Pdgfra-H2B-EGFP* transgenic mice (Hamilton et al., 2003) according to published isolation strategies (Kim et al., 2020). Cultured mesenchyme changed morphology in response to TGFB1-treatment, aggregating into clusters of cells in a dose-dependent response to TGFB1 (**Figure 5B-C**), confirming their ability to respond to the ligand. To identify the role of TGFB1 signaling in the interactions between the epithelium and mesenchyme, we co-cultured these cell types and manipulated TGFB-signaling (**Figure 5D**). Organoids were cultured for 3 days in 3D matrix bubbles and then floated above mesenchyme monolayers for 2 days before collecting the epithelium for qRT-PCR. Pre-treating the mesenchyme cultures with TGFB1 influenced the subsequent co-cultures, with a dose-dependent induction of transcripts associated with regeneration in the epithelium (**Figure 5E**).

**Figure 5.**
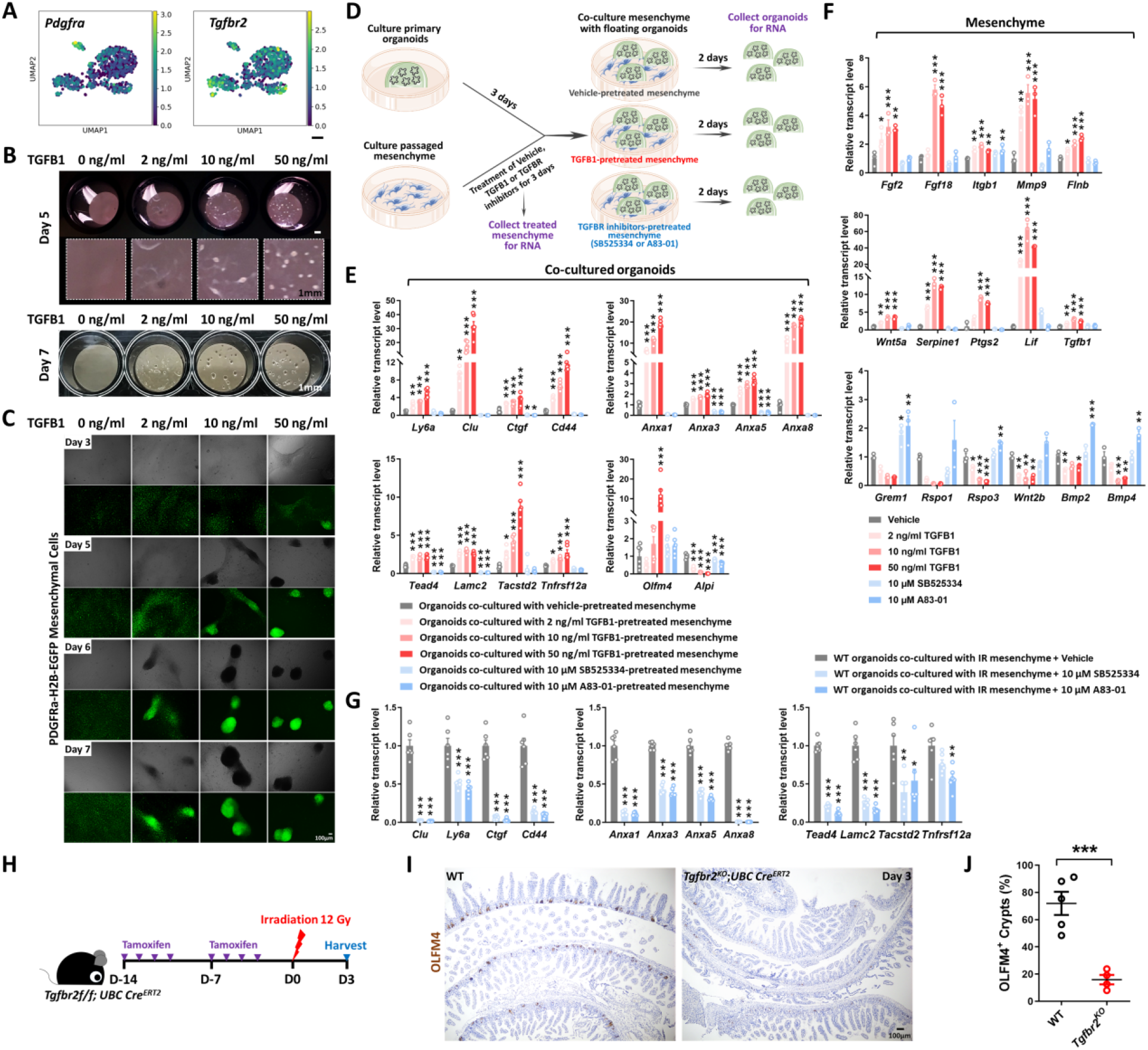
TGFB1-treated mesenchyme promotes fetal-like conversion of intestinal organoids. **(A)** UMAP indicates *Pdgfra*-positive mesenchymal cells express *Tgfbr2. Pdgfra* positive mesenchymal cells were subset from the scRNA-seq data set featured in **Figure 2** (GSE165318). **(B-C)** TGFB1-induces aggregation of *Pdgfra*-positive mesenchymal cells in a dose- and time-dependent manner (n=3 independent experiments, passaged mesenchyme). **(D-F)** TGFB1-treated mesenchyme induces fetal-like gene signatures in intestinal organoids upon co-culture. **(D)** Schematic of experimental design of co-culture. Passaged intestinal mesenchyme cells were pre-treated with vehicle, TGFB1 or TGFBR inhibitors for 3 days, and then co-cultured as overlaid matrigel bubbles containing primary organoids at day 3 post isolation. After 2 days of co-culture, organoids were collected in their matrix bubbles for qRT-PCR (n=6 independent organoid cultures with 2 different cell densities of mesenchyme). TGFB1 was removed during co-culture and only used for pre-treatment. TGFBR inhibitors were either kept **(E)** or removed (**Figure S5E**) in co-culture. **(F)** Mesenchyme cells pre-treated with vehicle, TGFB1 or TGFBR inhibitors for 3 days were also collected for qRT-PCR (n=3 independent mesenchyme cultures). **(G)** Presence of TGFBR inhibitors suppresses fetal-like conversion of intestinal organoids co-cultured with mesenchyme isolated from mice 3 days post-irradiation (n=6 independent organoid cultures with 2 different cell lines of mesenchyme). All the qRT-PCR data are presented as mean ± SEM. Transcript levels relative to vehicle control, and statistical comparisons were performed using one-way ANOVA followed by Dunnett’s post at *P* < 0.001***, *P* < 0.01** or *P* < 0.05*. **(H-J)** *Tgfbr2* knockout via *UBC-Cre*^*ERT2*^ restricts regeneration after irradiation. 5-week old mice were treated with tamoxifen to inactivate *Tgfbr2* in the whole body. Intestine was collected 3 days post-IR and scored for regenerative foci using OLFM4 immunostaining (n=4-5 biological replicates, duodenum, Student’s t-test at *P* < 0.001***). **(H)** Schematic of experimental design. **(I)** Representative images. **(J)** Quantification.

Conversely, pre-treatment with inhibitors of TGFB receptors suppressed the same markers of regeneration in the epithelium (**Figure 5E**). In response to TGFB1 pre-treatment, mesenchymal cells exhibited higher levels of transcripts expected to promote regeneration/wound healing such as *Ptgs2, Wnt5a*, and *Lif* (Miyoshi et al., 2012; Roulis et al., 2020; Wang et al., 2020); lower levels of transcripts that promote homeostatic epithelial growth, such as *Grem1* and *Rspo3* (Goto et al., 2022; McCarthy et al., 2020) were observed in response to TGFB1 treatment (**Figure 5F**), suggesting that TGFB1 reshapes the signaling environment to favor regenerative growth. Similar changes in transcriptional profiles were observed *in vivo* within the *Pdgfra*-expressing populations in response to IR (**Figure S5A**), and mesenchyme isolated from irradiated intestines expressed higher levels of transcripts for pro-regenerative growth ligands and reduced levels of ligands associated with homeostatic growth (**Figure S5B**). A pro-regenerative signature was also robustly reduced when mesenchymal cultures derived from irradiated mice were pre-treated with TGFB-inhibitors before epithelial overlay (**Figure 5G, Figure S5C-E**). Finally, to define the role of TGFB in both mesenchyme and epithelial cells in response to IR *in vivo*, we utilized *Tgfbr2*^*f/f*^;*UBC-Cre*^*ERT2*^ mice to inactivate the receptor across all cell populations. We found loss of *Tgfbr2* in mesenchyme and epithelial cells led to a dramatic reduction of regenerative crypts following IR (**Figure 5H-J**). The impaired regeneration is more pronounced in the *UBC-Cre*^*ERT2*^ model of *Tgfbr2* loss in both mesenchyme and epithelial cells than in epithelial-specific *Tgfbr2* knockout (**Figure 3I**). These experiments implicate stromal cells in TGFB-mediated intestinal regeneration.

### TGFB1 induces a YAP-SOX9 regenerative circuit

To better appreciate the epithelial response to TGFB1, we conducted ATAC-seq and defined 1,647 genomic regions that gained chromatin accessibility in organoids treated with TGFB1 (**Figure 6A, Figure S6A-C, Supplementary Table 1**, Diffbind FDR < 0.01). Examples of regeneration marker genes with increased chromatin accessibility in response to TGFB1 treatment include *Anxa1, Cd44* and *Wnt5a* (**Figure 6B**). Regions of the genome with increased accessibility were enriched in transcription factor motifs known to bind SOX, TEAD, and SMAD families of transcription factors (**Figure 6C**, HOMER). By contrast, the 3,900 genomics regions more accessible in the vehicle-treated condition are enriched in transcription factor binding motifs that are associated with function of the homeostatic intestinal epithelium (**Figure 6C**). This chromatin accessibility analysis suggests that Hippo-TEAD, SOX9, and TGFB-SMAD signaling activity or expression are elevated in response to TGFB1 treatment. Notably, SOX9 protein levels are induced after IR of the intestine and SOX9-expressing cells coincide with cells expressing the regenerative factor YAP (**Figure 6D, Figure S6D-E**). In scRNA-seq data re-analyzed from **Figure 3B** to focus on epithelial cells, *Sox9* is elevated in response to IR, and corresponds to elevation of genes involved in YAP signaling and organization of the extracellular matrix (**Figure 6E-G**). Importantly, *Sox9* is co-expressed in cells producing *Tgfbr2*, suggesting a potentially direct connection between TGFB1 and Sox9 regulation (**Figure 6H**). In mice treated with clodronate liposomes to deplete monocyte/macrophages, SOX9 protein and transcript levels were notably reduced in the crypt domain at 3 days post-IR compared to controls (**Figure 6I, Figure S6F**).

**Figure 6.**
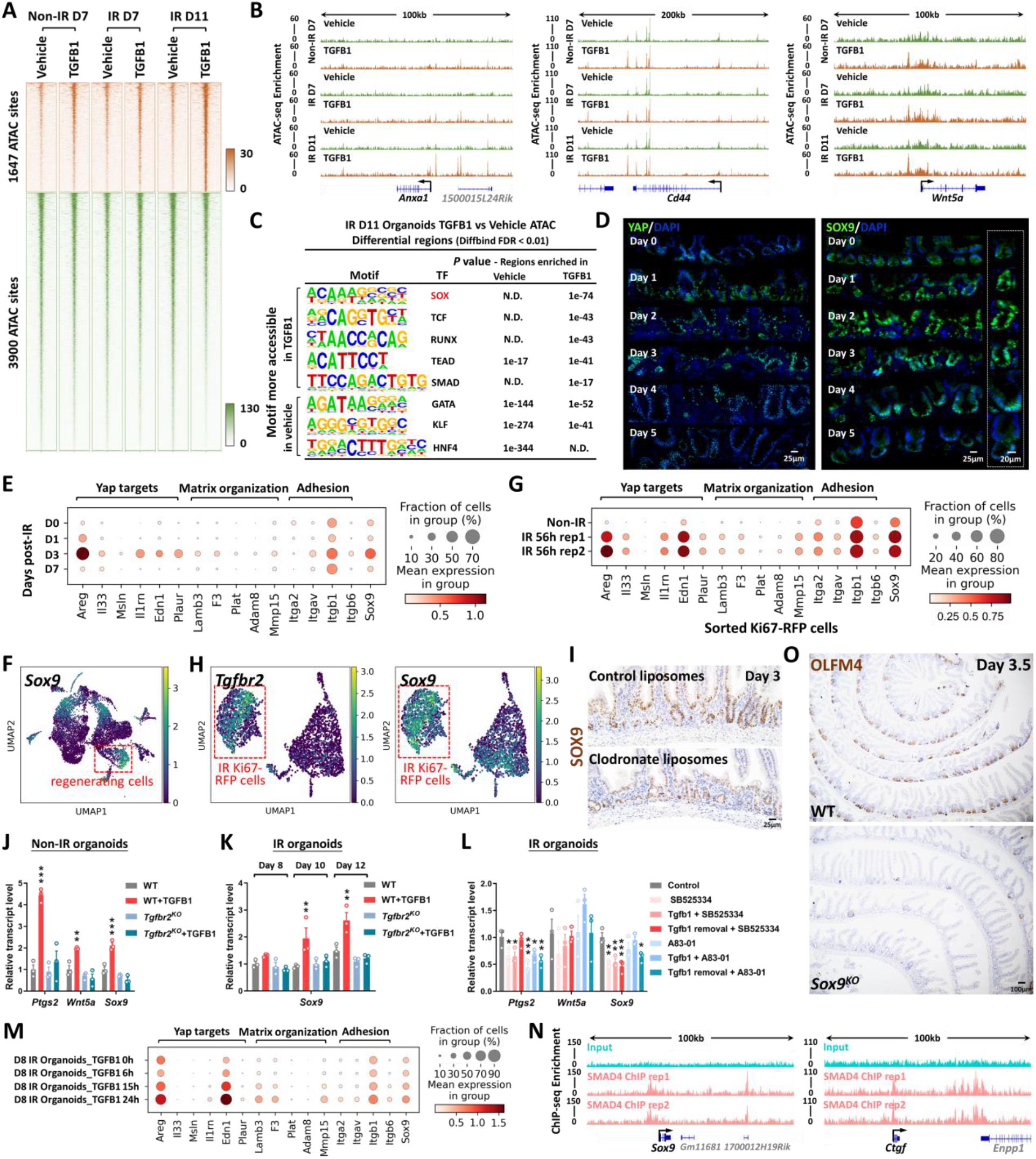
YAP-SOX9 circuit responds to TGFB1-induced open chromatin and epithelial regeneration in the intestine. **(A)** TGFB1-induced accessible chromatin identified using ATAC-seq Day 11 TGFB1-treated vs. vehicle-treated organoids (Diffbind FDR < 0.01, n=3 independent cultures). The experimental design is the same as for bulk RNA-seq and shown in **Figure 4A. (B)** Examples of genes harboring TGFB1-induced open chromatin visualized using IGV. **(C)** HOMER *de novo* DNA-motif enrichment analysis of ATAC-seq regions (Diffbind FDR < 0.01) shows that SOX, TCF, RUNX, TEAD and SMAD binding sequences are more prevalent in accessible regions of TGFB1-treated organoids, whereas GATA, KLF and HNF4 binding sequences are more prevalent in accessible regions of vehicle-treated organoids. N.D.: Not Detectable. **(D)** Immunofluorescence staining of YAP and SOX9 across a time course post-irradiation (representative of 3 biological replicates). Dot plots **(E)** and UMAP **(F)** reveal that YAP-related gene signatures and *Sox9* are elevated during regeneration post-irradiation, as evidenced by scRNA-seq (subset of intestinal epithelial cells, GSE165318). **(G)** scRNA-seq dot plots reveal that YAP related gene signatures and *Sox9* are also elevated in sorted Ki67-RFP positive cells after 56 hours of irradiation. **(H)** scRNA-seq UMAP reveals that *Tgfbr2*-positive cells express *Sox9* in sorted Ki67-RFP positive cells after 56 hours of irradiation. **(I)** Depletion of monocytes/macrophages (main cell sources of TGFB1 secretion) results in a downregulation of SOX9, as evidenced by immunostaining (representative of 3 biological replicates). **(J-L)** qRT-PCR indicates that TGFB1 induces expression of YAP related genes (*Ptgs2* and *Wnt5a*) and *Sox9* in WT organoids, and these effects are blocked in *Tgfbr2*^*KO*^ or TGFBR inhibitor-treated organoids. All the qRT-PCR data are presented as mean ± SEM (n=3 independent organoid cultures). Transcript levels are relative to WT, and statistical comparisons were performed using one-way ANOVA followed by Dunnett’s post at *P* < 0.001***, *P* < 0.01** or *P* < 0.05*. **(M)** Dot plots show that TGFB1 activates YAP related gene signatures and *Sox9* in a time-dependent manner (see experimental design in **Figure 4H**). **(N)** ChIP-seq shows that SMAD4 binds to gene loci of *Sox9* and *Ctgf* in mouse intestinal epithelium (GSE112946 (Chen et al., 2019)). **(O)** Loss of *Sox9* restricts regeneration after irradiation, as evidenced by OLFM4 immunostaining. Mice were treated with tamoxifen to inactivate *Sox9* in the intestinal epithelium 7 days before 12 Gy of irradiation. Intestine was collected 3.5 days post-IR (representative of 3 biological replicates).

Treatment of organoids with TGFB1 induced *Sox9* expression, and this induction was dependent upon the expression of *Tgfbr2*, could be blocked by TGFBR2-inhibitors, and increased with time after exposure to TGFB1 (**Figure 6J-M**). Organoid cells expressing *Sox9* in response to TGFB1 treatment (**Figure S6H**) were co-localized to cells expressing *Clu* (**Figure 4K**). ChIP-seq data indicate that SMAD4 can directly bind to the *Sox9* and *Ctgf* loci, suggesting a direct pathway between TGFB1 and the regenerative response (**Figure 6N**). SOX9 is critical for the regenerative response, as epithelial-specific inactivation of *Sox9* led to a severely reduced number of OLFM4^+^ regenerative crypts at 3 days post-IR (**Figure 6O**), similar to what has been reported by others (Roche et al., 2015). These data are consistent with a model in which TGFB1 exposure leads to direct transcriptional activation of *Sox9* and the YAP pathway, with subsequent activation of the regenerative response.

### Pre-treatment of organoid cultures with TGFB1 enhances engraftment efficiency into damaged colon

Epithelial transplants hold tremendous promise for cellular therapy to correct genetic disorders affecting the intestine or to accelerate healing of damaged mucosa. Given the importance of TGFB1 in promoting regenerative characteristics in the intestinal epithelium, we suspected that TGFB1 pre-treatment of epithelial organoid cultures could improve transplant efficiency. Since pre-irradiating organoids would not be conducive to use in the clinic, we revisited the strategy of treating non-irradiated organoids with TGFB1 and monitored the response using RNA-seq and qRT-PCR. We found TGFB1 induced fetal/regenerative gene expression is independent of IR-induced damage (**Figure 7A-D, Figure S7A**), and conserved in human organoids (**Figure S7B-E**). Loss of *Tgfbr2* suppresses fetal/regenerative gene expression in organoids (**Figure S7F**). Given a robust induction of regenerative marker gene expression in response to TGFB1 treatment in non-IR organoids (**Figure 7A-D**), we next assayed the effects of TGFB1 pre-treatment of organoids in a well-developed engraftment assay in which the host epithelium is damaged through exposing animals to Dextran Sulfate Sodium (DSS, (Watanabe et al., 2022)). Due to a variable response to DSS between mice, we conducted the assay as a competition experiment in which organoids treated with Vehicle or TGFB1 would be co-transplanted via enema. Organoids were collected from transgenic mice expressing fluorescent reporters and could be traced by immunofluorescence (**Figure 7E-F, Figure S7G**). Excitingly, TGFB1-treated organoids were significantly more likely to colonize the host colon than vehicle treated controls, as visualized by immunofluorescence (**Figure 7G, Figure S7H**). Engraftment, quantified either by the size of the individual graft or the average size of the organoid graft per mouse was significantly enhanced in the TGFB1-treated condition (**Figure 7H-I**). These data reflect the promise of TGFB1 pre-treatment in supporting the use of epithelial cultures for cellular therapy in the GI tract.

**Figure 7.**
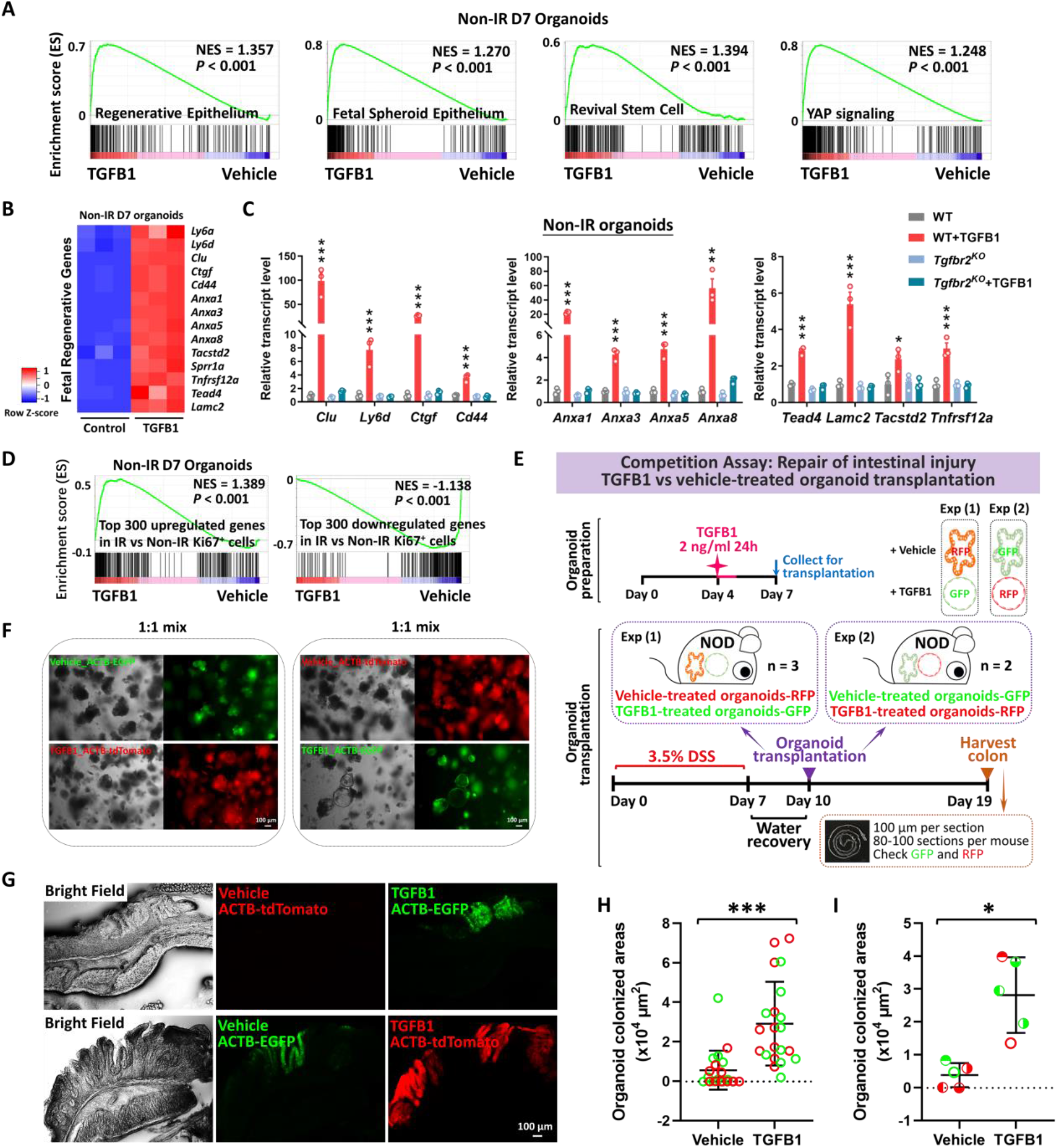
Transplantation of TGFB1-treated organoids enhances engraftment into DSS-treated mice. **(A)** Gene signatures of regenerative epithelium, fetal spheroids, revival stem cells and YAP signaling (Ayyaz et al., 2019; Gregorieff et al., 2015; Mustata et al., 2013; Wang et al., 2019; Yui et al., 2018) are each elevated post-TGFB1 treatment, as assayed by GSEA in non-irradiated conditions (n=3 independent organoid cultures, Kolmogorov-Smirnov test, *P* < 0.001). **(B)** Heatmaps display that RNA-seq expression levels of fetal/regenerative genes are highly expressed upon TGFB1 treatment compared to the vehicle controls (n=3 independent organoid cultures). Schematic of experimental design for bulk RNA-seq for panels A-B is depicted in **Figure S7A. (C)** For non-IR organoids, to deplete *Tgfbr2*, the primary organoids were treated with 1 μM tamoxifen for 12 hours on Day 3, followed with TGFB1 (2 ng/ml) treatment on Day 6. Organoids were collected 24 hours after TGFB1 treatment. Transcript levels are relative to WT; statistical comparisons were performed using one-way ANOVA followed by Dunnett’s post at *P* < 0.001***, *P* < 0.01** or *P* < 0.05* (n=3 independent organoid cultures). **(D)** GSEA reveals that genes upregulated or downregulated upon TGFB1 treatment strongly correlate with transcriptional changes in Ki67-RFP cells from the intestine of mice with irradiation vs. non-irradiation, respectively, as described in **Figure 1** (Kolmogorov-Smirnov test, *P* < 0.001, n=3 independent organoid cultures). **(E)** Experimental design for organoid transplantation assay, to determine the ability of TGFB1 to prime organoids in culture prior to transplantation for engrafting into damaged colonic tissue. Transgenic organoid lines were used to later help visualize transplants. Organoids were treated with either vehicle or with TGFB1 to induce regenerative properties. To induce epithelial damage in the mouse intestine, 3.5% DSS was prepared in drinking water and fed to NOD mice for 7 days. After a period of water recovery, the treated and control organoids were mixed 1:1 and used for enema-based transplant. Either vehicle-treated organoids with RFP were mixed with TGFB1-treated organoids with GFP; or vehicle-treated organoids with GFP were mixed with TGFB1-treated organoids with RFP. Organoid mixtures were transferred into DSS-treated mice on Day 10. Colon tissues were collected on Day 19, and cryosections were prepared for checking GFP or RFP under fluorescence microscope. **(F)** Representative images of organoids used for transplantation. **(G-I)** Representative images and quantification of transplant efficiency upon TGFB1 pre-treatment. **(G)** Fluorescent micrographs demonstrating transgenic organoid grafts into mice. **(H)** The size of grafts observed. **(I)** The average area of organoid grafts per mouse. Color indicates whether transplanted organoids derived from red or green fluorescent lines. Symbol type represents a single mouse used in the competition assay (n=21 grafts from 5 mice, Student’s t-test at *P* < 0.001*** or *P* < 0.05*).

## DISCUSSION

Radiotherapy, chemotherapy, inflammatory bowel disease, and other maladies of the gut induce extensive damage of the intestinal epithelial lining. These conditions could be alleviated by transplant of healthy epithelial cells. Genetic disorders affecting intestinal epithelial cell functions, such as microvillus inclusion disorders or congenital diarrhea or nutrient transporter deficiencies could also be corrected by replacement of defective epithelium with genetically corrected epithelial transplants. In this study, we discover a critical role for TGFB1 in promoting intestinal regeneration. TGFB signaling drives fibrosis in many organs including the intestine (Clouthier et al., 1997; Roberts et al., 2003; Sato et al., 2003; Sime et al., 1997; Vallance et al., 2005; Zanninelli et al., 2006; Zhao et al., 2002); however, our strategy of ex-vivo treatment could avoid tissue fibrosis. We propose a novel application of these findings to enhance intestinal engraftment by using TGFB1 ligands to induce a fetal-like regenerative-state in intestinal organoids. We demonstrate that TGFB1-treated organoids support more robust tissue engraftment in a mouse model of ulcerative colitis.

Mechanistically, we demonstrate pleiotropic functions for TGFB1 during regeneration. These TGFB1 functions include the induction of *Clu*^+^ cells, the promotion of fetal/regenerative gene signatures, the stimulation of mesenchymal cells to secrete pro-regenerative ligands, and the subsequent induction of a YAP-SOX9 circuit in the intestinal epithelium. *Clu*^+^ cells are very rare in homoeostatic intestine, but are activated in damaged intestine and can rapidly expand to reconstitute *Lgr5*^+^ ISCs and promote regeneration of the intestinal epithelium (Ayyaz et al., 2019). These *Clu*^+^ cells undergo a YAP-dependent transient expansion upon intestinal injury (Ayyaz et al., 2019). A fetal-like reversion of the regenerative epithelium and re-initiation of a fetal-like developmental transcriptional program were observed during intestinal regeneration after injury caused by DSS-induced colitis or helminth infection (Nusse et al., 2018; Yui et al., 2018). Upon injury, YAP transiently reprograms *Lgr5*^+^ ISCs by inducing a regenerative program while suppressing the Wnt-dependent homeostatic program (Gregorieff et al., 2015). We bring insight into the function of TGFB1 in the regeneration process, and suggest a clear advantage for TGFB1-treated organoids to engraft in DSS-damaged intestines compared to canonically-treated intestines.

Recent studies suggest mesenchymal cells as important sources of niche signals (Aoki et al., 2016; Degirmenci et al., 2018; Greicius et al., 2018; McCarthy et al., 2020; Shoshkes-Carmel et al., 2018; Stzepourginski et al., 2017). We reveal that TGFB1 pushes mesenchymal cells from homeostatic RSPO/Grem1 signals to regeneration/wound healing signals. Elevated levels of *Ptgs2* and *Wnt5a* were observed in TGFB1-treated mesenchymal cells. *Ptgs2*-expressing fibroblasts process arachidonic acid into prostaglandin E2 (PGE2), which enhances YAP activity through the prostaglandin EP4 receptor and drives expansion of Sca-1^+^ reserve-like stem cells (Roulis et al., 2020). PGE2 triggers cell fate plasticity by promoting a switch from differentiated enterocytes to wound-associated epithelial (WAE) cells (Miyoshi et al., 2017). Additionally, Wnt5a, a noncanonical Wnt ligand, potentiates TGFB signaling and is required for crypt regeneration through the WAE cells (Miyoshi et al., 2012). Interestingly, *Trem2*-expressing macrophages associate with the WAE layer (Seno et al., 2009), and WNT-producing macrophages enhance epithelial regeneration following tissue damage (Saha et al., 2016). While previous studies have demonstrated TGFB1 production by circulating macrophages following phagocytosis of dying cells (Fadok et al., 1998), here we demonstrate that macrophages are the primary TGFB1 source in intestinal regeneration, and the TGFB1 is necessary for full regeneration and sufficient to induce the epithelial regenerative state. Integration of our *in vivo* findings and the growing knowledge of intestinal regeneration with *in vitro* approaches to induce regenerative epithelium should accelerate progress towards cellular therapies.

## METHODS

### Mice and treatment

The *Villin-Cre*^*ERT2*^ transgene (el Marjou et al., 2004), *UBC-Cre*^*ERT2*^ transgene (Ruzankina et al., 2007), *Tgfbr2*^*f/f*^ (Leveen et al., 2002), *Smad4*^*f/f*^ (Yang et al., 2002), *and Sox9*^*f/f*^ (Akiyama et al., 2002) alleles were integrated to generate the conditional compound-mutants and controls. For *Villin-Cre*^*ERT2*^*;Smad4*^*f/f*^, mice (8-12 weeks old) were treated with tamoxifen (Sigma T5648) at 50 mg/kg/day for 4 consecutive days by intraperitoneal injection. For *Villin-Cre*^*ERT2*^*;Tgfbr2*^*f/f*^ and *Villin-Cre*^*ERT2*^*;Sox9*^*f/f*^, mice (8-12 weeks old) were treated with tamoxifen at 100 mg/kg/day for 4 consecutive days. The knockout efficiency varied in *Sox9*^*KO*^, and only the mice with more than 75% of Sox9 depletion were used for downstream analysis. For *UBC-Cre*^*ERT2*^*;Tgfbr2*^*f/f*^, 5-week-old mice were treated with tamoxifen at 100 mg/kg/day for 4 consecutive days at week 5. The same treatment was repeated at week 6. C57BL/6, *Mki67tm1*.*1Cle/J* (also known as *Ki67-RFP*) (Basak et al., 2014), *PDGFRa-H2B-EGFP* (Hamilton et al., 2003), *ACTB-EGFP* (Okabe et al., 1997), *ROSA26mT/mG* (ACTB-tdTomato,-EGFP) (Muzumdar et al., 2007), and NOD SCID (Blunt et al., 1995) mice were also used in this study. KAPA Mouse Genotyping Kits (Kapa Biosystems, KK7352) were used to identify genotypes of mice.

For IR treatment, mice were subjected to 12 Gy whole-body IR with a 137 Cs γ-source irradiator at a dose rate of 60 or 85 cGy/minute. Experiments were analyzed within groups exposed to the same irradiator treatment. For the BrdU pulse-chase experiment, mice were injected with 1 mg BrdU by intraperitoneal injection. To deplete monocytes/macrophages, C57BL/6 mice were treated with clodronate-containing liposomes (FormuMax Scientific, SKU: F70101C-NC-10) by intraperitoneal injection (2 treatments of 200 μl, 72 hours pre- and day of IR). Treatment with control liposomes (same treatment course and dose) was performed as a control. To neutralize TGFB, C57BL/6 mice were treated with TGFB antibody (1D11, MA5-23795, Invitrogen) by intraperitoneal injection (250 μg per dose, two doses, on the day right after IR and 1 day after IR). Mouse IgG1 Isotype (MAB002, R&D) or vehicle were used as control. All mouse protocols and experiments were approved by the Rutgers Institutional Animal Care and Use Committee.

### Histology and immunostaining

Freshly harvested intestinal tissues were fixed overnight with 4% paraformaldehyde at 4°C, and then washed with PBS. For paraffin embedding, tissues were then dehydrated through ascending alcohols and processed with xylene prior to embedding. For cryo-embedding, tissues were then processed with 15% sucrose and 30% sucrose until tissues sunk prior to freezing in OCT compound (Tissue-Tek 4583). 5 μm-thick paraffin sections and 10 μm-thick cryosections were used for immunohistochemistry and immunofluorescence using standard procedures, respectively. Hematoxylin (VWR, 95057-858) and eosin (Sigma, HT110180) staining was performed using standard procedures. Immunohistochemistry was performed using primary antibodies against Ki67 (Abcam ab16667, 1:300), OLFM4 (Cell Signaling 39141, 1:2500), CD44 (BD 558739, 1:300), BrdU (Bio-Rad MCA2060, 1:500), F4/80 (Cell Signaling 70076, 1:500), and SOX9 (Cell Signaling 82630, 1:600). After incubating with secondary antibody and the Vectastain ABC HRP Kit (Vector Labs), slides were developed using 0.05% DAB (Amresco 0430) and 0.015% hydrogen peroxide in 0.1 M Tris, and then counterstained with hematoxylin. The slides were mounted and viewed on a Nikon Eclipse E800 microscope. Images were photographed with a Lumenera INFINITY3 camera and infinity capture imaging software (v6.5.6). A Zeiss Observer Z1 microscope was used to image the immunofluorescence staining of YAP (Cell Signaling 4912, 1:100), SOX9 (Cell Signaling 82630, 1:100) and DAPI (Biotium 40043, 1:5000). ImageJ and Adobe Photoshop were used to adjust contrast and brightness. When adjustments of sharpness, contrast, or brightness were made, they were applied uniformly across comparative images.

### Intestinal crypt isolation

Freshly harvested intestine was flushed with cold PBS, opened longitudinally, cut into 1 cm pieces, and then rotated in 3 mM EDTA in PBS at 4 °C for 5 minutes, 10 minutes and 40 minutes (refreshing EDTA/PBS every time). The tissue was then vigorously shaken to release the epithelium, and crypts passed through a 70-μm cell strainer. Cells were pelleted by centrifugation at 200 g for 3 minutes at 4 °C and then washed with cold PBS. Cell pellets were used for organoid culture and single cell dissociation for ATAC-seq as described in later sections.

### Organoid culture and treatment

Primary crypt-derived organoids were isolated from proximal half of mouse small intestine and cultured in Cultrex® reduced growth factor basement membrane matrix, Type R1 (Trevigen) according to established methods (Sato et al., 2009). Organoid medium was changed every 2 days. The organoids were treated with 1 μM tamoxifen dissolved in ethanol for 12 h. Vehicle-treated organoids served as a control. Tamoxifen was added into culture medium of organoids on Day 3 after seeding. Recombinant TGFB1 (Peprotech 100-21) or 10 μM TGFB receptor inhibitors including SB525334 (Selleckchem S1476) and A83-01 (Tocris 2939) were prepared according to the supplier’s instructions, and vehicle controls were used in organoid culture. For IR treatment, organoids on Day 4 after seeding were subjected to 4 Gy IR with a 137 Cs γ-source irradiator at a dose rate of 60 or 85 cGy/minute. 1-2 ng/ml TGFB1 was added on Day 6 or Day 7 after seeding. Details of each experimental design can be found in figure schematics and figure legends. TGFB1 activity was tested before each experiment and the dose of TGFB1 was adjusted by its ability to induce spherical organoids, targeting at ∼60-80% of spherical organoids after irradiation in this study, according to the decline in TGFB1 activity over time in storage as determined (**Supplementary Table 2**).

Human duodenal organoid lines were cultured in Corning Matrigel (REF 356231) with L-WRN complete medium, which contains 50% L-WRN conditioned medium. Every 220 μL Matrigel stock was supplemented with 59 μL cold L-WRN complete medium, plus an additional 0.6 μL 2.5 mM Y27632, and 5.5 μL 100 μM CHIR99021. Conditioned media from L-WRN cells containing Wnt3a, Rspondin3, and Noggin was mixed 1:1 with 2x Basal media comprised of 43.8 mL advanced DMEM/F-12, 1 mL 200 mM GlutaMAX, 1 mL 1 M HEPES, 1 mL N-2 supplement, 2 mL B-27 supplement, 200 μL 0.5 M N-Acetyl-L-cysteine, and 1 mL penicillin/streptomycin. To make L-WRN complete medium, the mixed media mentioned above was further supplemented with 50 ng/mL human EGF, 100 μg/mL primocin, 2.5 μM CHIR99021 and 10 μM Y27632. To investigate the conserved function of TGFB1 in human, 2 ng/ml recombinant TGFB1 (Peprotech 100-21) and vehicle controls were used in human duodenal organoid culture for 24 hours. The human organoids were collected 24 hours or 72 hours after TGFB1 initiation.

### Single cell dissociation of organoids for ATAC-seq and scRNA-seq

Mouse organoids were cultured and treated as mentioned above. Primary cultured organoids were collected by removing Matrigel using cold PBS. Matrigel droplets containing organoids were disrupted. Organoids were pelleted by centrifugation at 300 g for 3 minutes at 4 °C and washed with cold PBS. After removing the PBS and Matrigel, organoids were resuspended in 1 mL pre-warmed (37 °C) TrypLE, and rotated at 37 °C for 15 to 30 minutes until organoids were dissociated into single cells (confirmed via microscope). Cells were pelleted at 300 g for 3 minutes at 4 °C, and washed with cold PBS. Cells were passed through SP Bel-Art Flowmi 40 μm cell strainer and collected into a new protein LoBind tube. 30,000 cells were prepared as mentioned above and used for ATAC-seq as described previously (Buenrostro et al., 2013; Buenrostro et al., 2015) with slight modifications. Briefly, cells were centrifuged at 500 g for 5 minutes at 4°C and resuspended in ice-cold lysis buffer (10 mM Tris, pH 7.4, 10 mM NaCl, 3 mM MgCl2, and 0.1% NP-40). Cells were then centrifuged at 500 g for 10 minutes at 4°C. The isolated nuclear pellets were incubated with a 50 μl reaction of Tn5 Transposase (Illumina Tagment DNA Enzyme and Buffer Large Kit 20034198) for 30 minutes at 37°C. The transposed chromatin was purified with MinElute PCR purification kit (QIAGEN REF 28004), and PCR was amplified with high-fidelity 2x PCR Master Mix (New England Biolabs M0541). One-third of the maximum fluorescent intensity during a qPCR trial run was used to determine the additional cycles for library prep. The PCR amplified libraries were purified and sent to Novogene America for sequencing.

Cells used for scRNA-seq were prepared according to PIPseq Milli 3’ Single Cell Capture and Lysis User Guide. Briefly, a cell suspension at a concentration of 1000 live cells/μL was prepared in Cell Suspension Buffer provided from the PIPseq™ T2 3’ Single Cell Capture and Lysis Kit (v2.1, Fluent BioSciences). 5000 cells (total 5 μL) were added into Pre-templated Instant Partitions (PIPs). Stable emulsions carrying captured mRNA were sent to Fluent Biosciences for downstream processing. A visual inspection was performed at sample receipt to assess the emulsion quality before initiating downstream sample processing. Satisfactory samples were carried through mRNA isolation, cDNA generation, cDNA amplifications, sequencing ready library preparation, library pooling, and sequencing according to the PIPseq™ T2 Single Cell Sequencing Kit (v2.1) specifications. cDNA and library quality were assessed by using ThermoFisher Qubit 4 Fluorometer and Agilent 4200 TapeStation System. cDNA libraries were sequenced on an Illumina NextSeq 2000 instrument to a minimum sequencing depth of 50,000 reads per cell.

### Fluorescence-Activated Cell Sorting (FACS) for scRNA-seq

Crypts were isolated from proximal half of small intestine of Mki67tm1.1Cle/J mice after 56 hours post-irradiation and their non-IR littermate control, as described in the previous section. 100-μm cell strainer was used in IR samples, as crypts were expanded during regeneration after IR. Crypts were pelleted by centrifugation at 300 g for 3 minutes at 4 °C and then washed with cold PBS. To dissociate single cells for FACS, isolated crypts were further rotated with 5U/ml dispase (Stem Cell 07913) and 200 U/ml of DNase I (Sigma D4513) at 37 °C for 30 minutes, and were then washed twice with 1% BSA/PBS, and filtered with a 40-μm cell strainer. Cells were prepared in 1% BSA/PBS with 200 U/ml of DNase I for sorting. Ki67-RFP^+^ DAPI^-^ cells from mice 56 hours post-IR and non-IR condition were detected and sorted with Beckman Coulter Astrios EQ High Speed Cell Sorter, respectively. Dead cells were eliminated using 0.5 μg/ml DAPI. Kaluza analysis 2.1.3 software was used for FACS data analysis (**Supplementary Table 3**). Total 5500 cells were sorted and concentrated into 5 μL for initiating the PIPseq pipeline (PIPseq™ T2 3’ Single Cell Capture and Lysis Kit v2.1, Fluent BioSciences). Cells were captured and lysed according to the manufacturer’s instructions. Samples were sent to Fluent BioSciences for downstream processing as described above.

### Isolation and culture of intestinal mesenchymal cells

Mesenchymal cells were isolated from proximal half of small intestine from C57BL/6 or PDGFRa-H2B-EGFP mice as previously described (Kim et al., 2020) with slight modifications. Freshly harvested intestine was flushed with cold PBS, opened longitudinally, cut into 2 cm pieces, and then rotated in 30 mL pre-digestion buffer (pre-warmed, HBSS containing 10% FBS, 10 mM HEPES and 5 mM EDTA) at 37°C for 20 minutes twice (refresh pre-digestion buffer every time). After washing tissues in wash buffer (HBSS containing 2% FBS, 10 mM HEPES), the tissues were further transferred into 20 mL of pre-warmed digestion buffer (RPMI medium containing 10% FBS, 1% P/S, 15 mM HEPES, 25 U/mL of collagenase IV (Worthington LS004186), 100 U/ml of DNase I (Sigma D4513), 0.3 g/100 mL of Dispase II (Gibco 17105041)) and rotated at 37°C for 30 minutes. After vortexing the cell solution intensely for 20 sec every 10 minutes, the cell solution was passed through a 40 μm cell strainer. Cells were pelleted by centrifugation at 400 g for 5 minutes at 4 °C and then resuspended in RPMI medium containing 10% FBS. Remaining tissues were incubated with 20 mL of fresh digestion buffer, and the above steps were repeated. Cells were combined, seeded at a desired density (see below), and cultured in Advanced DMEM/F12 (Gibco 12634-010) medium, containing 10% FBS (Gibco 26140-095), 1% penicillin and streptomycin (Invitrogen 15140-122), 1% HEPES (Gibco 15630-080) and 1% Glutamax (Gibco 35050-061).

### Co-culture of intestinal mesenchymal cells and organoids

To set up a co-culture system, 3 intact 25ul Matrigel droplets were gently taken out from a cultured primary organoid plate 3 days after seeding. These Matrigel droplets containing organoids were transferred and floated above the cultured mesenchyme for 2 days. Co-cultured organoids were then collected for RNA extraction. Primary or passaged mesenchymal cells were cultured in 12-well plates. Primary mesenchymal cells isolated from IR and non-IR C57BL/6 mice were seeded at three densities, including 4×10^5^ cells, 10^6^ cells, 2×10^6^ cells. The cultures were refreshed at Day 3 to remove any debris. Co-culture was initiated on Day 4. For passaged mesenchymal cells, cells were seeded at two densities, including 10^5^ cells and 2.5×10^5^ cells. These cells were pre-treated with vehicle, TGFB1 or TGFBR inhibitors for 3 days, starting at Day 1 after seeding. These pre-treated mesenchymal cells were then used for co-culture. The detailed schematics of co-culture experiments are shown in **Figure 5D** and **Figure S5C**.

### RNA extraction, bulk RNA-seq, and quantitative reverse transcription polymerase chain reaction (qRT-PCR)

For tissues, 2 cm of mouse duodenum was homogenized in Trizol (Invitrogen 15596018), and processed for RNA extraction according to the manufacturer’s instructions. For cultured organoids or mesenchymal cells, QIAGEN RNeasy Micro Kit was used to extract RNA according to the manufacturer’s instructions.

For bulk RNA-seq, RNA samples were prepared and sent to BGI Americas. For qRT-PCR, cDNA was synthesized from total RNA with Oligo(dT)20 primers using SuperScript III First-Strand Synthesis SuperMix (Invitrogen 18080-400). qRT-PCR analysis was performed using gene-specific primers and SYBR Green PCR Master Mix (Applied Biosystems, 4309155). The sequences of the primers used are available upon request. The 2^-ΔΔCt^ method was applied to calculate the fold change of relative transcript levels, and *Hprt* was used for normalization.

### Protein extraction and western blot

2 cm of mouse duodenum was homogenized in RIPA buffer (50 mM Tris-HCl pH 8.0, 150 mM NaCl, 1% NP-40, 0.5% Na-deoxycholate, 0.1% SDS, protease inhibitor cocktails, and phosphatase inhibitors) and rotated at 4°C for 30 min for protein extraction. Protein concentration was determined by Pierce BCA Protein Assay Kit (Thermo). Immunodetection was performed using specific antibodies against p-SMAD3 (Cell signaling 9520, 1:1000), p-SMAD2/3 (Cell signaling 8828, 1:1000), SMAD2/3 (Cell signaling 8685, 1:1000), and β-actin (Abcam ab8227, 1:5000). The intensity of signal was quantified by ImageJ.

### Membrane-based antibody arrays

Protein lysates were extracted from 2 cm of duodenal fragment of mice after 3 days of irradiation, and non-IR mice were used as controls. The protein lysates (200 μg) were applied to Mouse Growth Factor Array C3 (RayBiotech, CODE: AAM-GF-3-4) according to the manufacturer’s instructions. Each growth factor was represented in duplicate on the membrane. Two independent experiments were performed to evaluate the expression level of various growth factors. The intensity of signal was quantified by ImageJ. The positive control was used to normalize the results from different membranes being compared.

### ELISA of TGFB1

Protein lysates were extracted from 1) 2 cm of duodenal tissues of mice after 3 days of irradiation, and non-IR mice were used as controls; 2) 2 cm of duodenal tissues of mice with clodronate liposomes, and mice treated with control liposomes were used as controls. Blood was collected from abdominal aorta of experimental mice mentioned above with heparinized syringes. Blood samples were centrifuged at 1500g for 15 minutes at 4 °C and the supernatant was collected as the plasma. To detect the relative abundance of TGFB1 in the intestine tissues and plasma, the protein lysates or plasma were applied to Mouse TGF beta 1 ELISA Kit (ab119557, abcam) according to the manufacturer’s instructions.

### RNAscope *in situ* hybridization and immunofluorescence

RNA localization was performed according to the RNAscope Multiplex Fluorescent Reagent Kit v2 Assay (ACD 323110) as previously described (Holloway et al., 2021). Briefly, mouse intestines were linearized and twisted into a swiss roll and prepared by 24-hour fixation in 10% normal buffered formalin at room temperature followed by dehydration through a methanol series and rehydration to 70% ethanol before paraffin infusion. Paraffin sections were cut at 5 μm, baked for one hour at 60°C, deparaffinized and then pretreated with hydrogen peroxide (ACD kit) for 10 minutes, antigen retrieval (ACD 322000) for 15 minutes and protease plus treatment (ACD kit) for 30 minutes. Probes were Mm-Tgfb1 (407751) and Mm-Tgfbr2-C2 (406241-C2) diluted 1:50 into the channel 1 probe. TSA fluorophores were diluted in multiplex TSA buffer (ACD 322809) Cy5 (Akoya TB-000203) at 1:2000, Cy3 (Akoya TB-000202) diluted 1:2500 and OPAL-488 (Perkin Elmer FP1168) at 1:2000. Following RNAscope, sections were co-stained with rabbit anti-F4/80 (Cell Signaling 70076, 1:100) and mouse-anti-Ecadherin (BD Transduction Labs 6101 1:500). Secondary antibodies were anti-Rabbit-750 (Sigma SAB4600373) and anti-mouse-594 (Jackson ImmunoLabs 715-585-150) both diluted at 1:500. Slides were mounted with Prolong Gold and imaged on an Olympus FV3000 confocal microscope.

### Organoid transplantation, imaging and quantification

Colitis was induced in NOD SCID female mice by administration of 3.5% Dextran sulfate sodium salt (DSS, Affymetrix J14489) in drinking water for 7 days followed by normal water for 3 days. Transplantation was performed 10 days after DSS initiation. The procedure was performed as previously described (Watanabe et al., 2022; Yui et al., 2018) with slight modifications. Donor crypts from ACTB-EGFP or ROSA26mT/mG (ACTB-tdTomato) mice were isolated from proximal half of small intestine and cultured in Cultrex® reduced growth factor basement membrane matrix, Type R1 (Trevigen) according to established methods (Sato et al., 2009). On Day 4 of primary culture, 2 ng/ml TGFB1 or vehicle was added into culture medium and removed 24 hours later. On Day 7 of primary culture, organoids were collected by removing Matrigel using cold PBS. To minimize variations, vehicle- and TGFB1-treated organoids were 1:1 mixed as follows and transferred to the same mouse: vehicle-treated organoids expressing RFP (ACTB-tdTomato) were mixed with TGFB1-treated organoids expressing GFP (ACTB-EGFP); or vehicle-treated organoids expressing GFP (ACTB-EGFP) was mixed with TGFB1-treated organoids expressing RFP (ACTB-tdTomato). In this competitive assay of repair of intestinal injury, approximately 600 organoids were suspended in 50 μl of 5% Matrigel in PBS for transplantation of each mouse.

Immune compromised NOD SCID female mice were anesthetized with isoflurane, and their colons were flushed using a gentle sterile PBS enema, followed by gentle massage of the abdomen to expel enema fluid and colon contents. The cell suspension mix of organoids was subsequently infused into the colonic lumen. The anus was sealed with surgical skin glue, which was removed after 6 hours. Colon tissues were collected after 9 days of organoid colonization. Colons were washed with PBS, opened longitudinally and pinned on a wax plate. After soaking in 4% paraformaldehyde for 10 minutes, swiss rolls were made for these colon tissues, and fixed for 2 hours with 10 ml of 4% paraformaldehyde at 4°C, and then washed with PBS. For cryo-embedding, tissues were then processed with 15% sucrose for 1 hour and 30% sucrose overnight at 4°C prior to freezing in OCT compound (Tissue-Tek 4583). The whole cryo block from each mouse was processed into sequential sectioning of 100 μm-thick cryosections, 80-100 sections per mouse. These sequential cryosections were imaged for GFP or RFP colonized regions of organoids under a Zeiss Observer Z1 microscope. ImageJ was used to quantify the organoid colonized areas from each grafts observed.

### Bioinformatics

For RNA-seq, raw sequencing reads (fastq) were first quality checked using fastQC (v0.11.3) and were further aligned to mouse (mm9) genomes using Tophat2 (v2.1.0) to generate bam files. Cuffquant (v2.2.1) was used to generate cxb files from bam files. Cuffnorm (v2.2.1) was performed to calculate FPKM values using quartile normalization. Cuffdiff (v2.2.1) was applied to identify differentially expressed genes between groups using quartile normalization and per-condition dispersion. Gene set enrichment analysis (GSEA v4.0.3) (Subramanian et al., 2005) was performed as described, and the gene signature lists used in this study were shown in **Supplementary Table 4**. Heatmapper (Babicki et al., 2016) was used to display relative transcript levels of genes of interest by using normalized FPKM values from Cuffnorm.

For scRNA-seq of organoid samples, prior to processing, a mouse STAR reference was generated using a mouse genome FASTA build from the Ensembl 102 release and GTF annotations from GENCODE (GRCm39 vM29 2022.04) using STAR’s genomeGenerate function. BCLs derived from Illumina sequencing of PIPseq samples were demultiplexed and output in FASTQ format using BCL-convert v4.0.3. Barcode whitelisting and error correction were performed using PIPseeker v0.55. Reads passing this stage were then aligned to the mouse reference genome using STARsolo (STAR v2.7.10a). An aligned, sorted BAM file was generated by STAR, which included required information for RNA velocity analysis. Cell calling was performed using PIPseeker’s transcript-count thresholding approach and sensitivity 3 was chosen for all samples, which most closely targeted the first inflection point after the knee point on the barcode-rank plot. For sorted Ki67-RFP^+^ DAPI^-^ cells, samples were processed in the same way, but using salmon v1.4 (Bala et al., 2022) instead of STAR. A reference genome was first built from the same sources as above using the salmon index command with a 19-bp k-mer size. The cell barcodes files from the filtered matrix, corresponding to the cell fraction, were then processed with Scanpy v1.9.1 (Wolf et al., 2018). RNA velocity was analyzed by Velocyto 0.17 (La Manno et al., 2018).

For ATAC-seq, paired-end ATAC-seq fastq file was quality checked using fastQC. ATAC-seq adapter sequences were removed from each read file using CutAdapt (v1.9.1) (Martin, 2011). Each read file was then aligned to the mouse mm9 genome using bowtie2. Picard (v2.18.27) was used to determine the median alignment size of each alignment bam file. Peak region bed files were called from each alignment bam file using MACS2 (v2.1.0). MACS2 was run with a “shift” distance of −0.5 times the median alignment size and an “extsize” distance equal to the median alignment size for each alignment bed file. The peak files were filtered against ENCODE blacklist, which is known to yield false ChIP-seq signals due to the inaccuracies of a particular genome assembly (Amemiya et al., 2019). BEDtools (Quinlan and Hall, 2010) was used to subtract the sites on the ENCODE blacklist. DiffBind (v2.4.7) (Stark and Brown, 2011) was used to identify differential signals of ATAC-seq between vehicle- or TGFB1-treated organoids, and FDR < 0.01 was used as the cutoff for significance. Haystack (v0.4.0) (Pinello et al., 2018) quantile normalized bigwigs were used to plot heatmaps using computeMatrix and plotHeatmap from deeptools (v2.4.2) (Ramirez et al., 2016). SitePro (v1.0.2) (Shin et al., 2009) was used to visualize the average signals of ATAC-seq in the desired genomic regions. Homer findMotifsGenome.pl (v4.8.3, homer *de novo* Results) (Heinz et al., 2010) was used to identify transcription factor motifs enriched at peaks. The Integrative Genomics Viewer (IGV 2.4.13) (Robinson et al., 2011) was used to visualize ATAC-seq bigwig tracks.

For ChIP-seq, FastQC (v0.11.3) was used to check the quality of raw sequencing reads (fastq), and bowtie2 (v2.2.6) was used to align the sequences to mouse (mm9) genomes and generate bam files. Model-based Analysis of ChIP-Seq (Zhang et al., 2008) (MACS 1.4.1) was used for peak calling and to generate bed files from aligned reads. The shiftsize parameter used in MACS was based on the fragment size of Pippin Prep library size selection. SMAD4 ChIP-seq of mouse duodenal epithelium are at a MACS p-value of 10^−5^. IGV 2.4.13 (Robinson et al., 2011) was used to visualize ChIP-seq bigwig tracks.

### Statistical analysis

The data is presented as mean ± SEM, and statistical comparisons were performed using one-way ANOVA followed by Dunnett’s post test with the GraphPad Prism version 8.0.2 or Student’s t-test at *P* < 0.001***, *P* < 0.01** or *P* < 0.05*. Other bioinformatics related statistical analysis was completed with the embedded statistics in each package, including MACS or MACS2 (Zhang et al., 2008), HOMER (Heinz et al., 2010), DiffBind (Stark and Brown, 2011), Cuffdiff (Trapnell et al., 2012), GSEA (Subramanian et al., 2005; Tamayo et al., 2016) and Scanpy (Wolf et al., 2018). *P* < 0.05 (95% confidence interval) was considered statistically significant.

### Data availability

All RNA-seq, scRNA-seq and ATAC-seq data of this study have been deposited in GEO (GSE222505). The following datasets from GEO were reanalyzed with our sequencing data: the accession number for the SMAD4 ChIP in mouse intestinal epithelium from our previous studies is GSE112946 (Chen et al., 2019). Bulk RNA-seq of GSE165157 (Qu et al., 2021) was used to analyze crypt transcriptome upon IR at days 0, 1, 2, 3 and 5. scRNA-seq of GSE117783 (Ayyaz et al., 2019) was used to compare normal crypts and irradiated crypt cells after 3 days of irradiation. scRNA-seq of GSE165318 was used to analyze duodenum/jejunum boundary samples collected at days 0, 1, 3, 7, and 14 after irradiation. scRNA-seq of GSE145866 (Sheng et al., 2020) was used to analyze sorted *Msi1*-GFP positive cells (irradiation-resistant) and their progeny cells upon irradiation.

## ACKNOWLEDGMENTS

M.P.V is supported by grants from the National Institutes of Health (NIH R01 DK126446 and R01 DK121915). M.P.V. and J.R.S are supported by the Intestinal Stem Cell Consortium from the National Institute of Diabetes and Digestive and Kidney Diseases (NIDDK) and National Institute of Allergy and Infectious Diseases (NIAID) of the NIH under grant number U01 DK103141. L.C. is supported by grants from Start-Up funding of Southeast University and National Natural Science Foundation of China 32270830. K.D.W. is supported by grant from NIH R01 DK121166. The authors acknowledge the Office of Advanced Research Computing (OARC) at Rutgers University for providing access to the Amarel cluster and associated research computing resources that have contributed to the results reported here. The research was also supported by flow cytometry/cell sorting core facility at Environmental and Occupational Health Sciences Institute (EOHSI) at Rutgers University. This work benefited from the Cancer Center Support Grant (CCSG, P30CA072720) from the National Cancer Institute.

## AUTHOR CONTRIBUTIONS

L.C. conceived and designed the study; performed benchwork, sequencing data processing, and bioinformatics; collected and analyzed the data; and wrote the manuscript. A.D., O.P.C. and J.W. contributed to benchwork. X.Q. and K.D.W. contributed to benchwork and data collection. A.O.P., W.H., and J.P.S. provided experimental materials or instruments. M.P.V. conceived and supervised the study, and wrote the manuscript.

## FIGURE AND FIGURE LEGENDS

**Figure S1.**
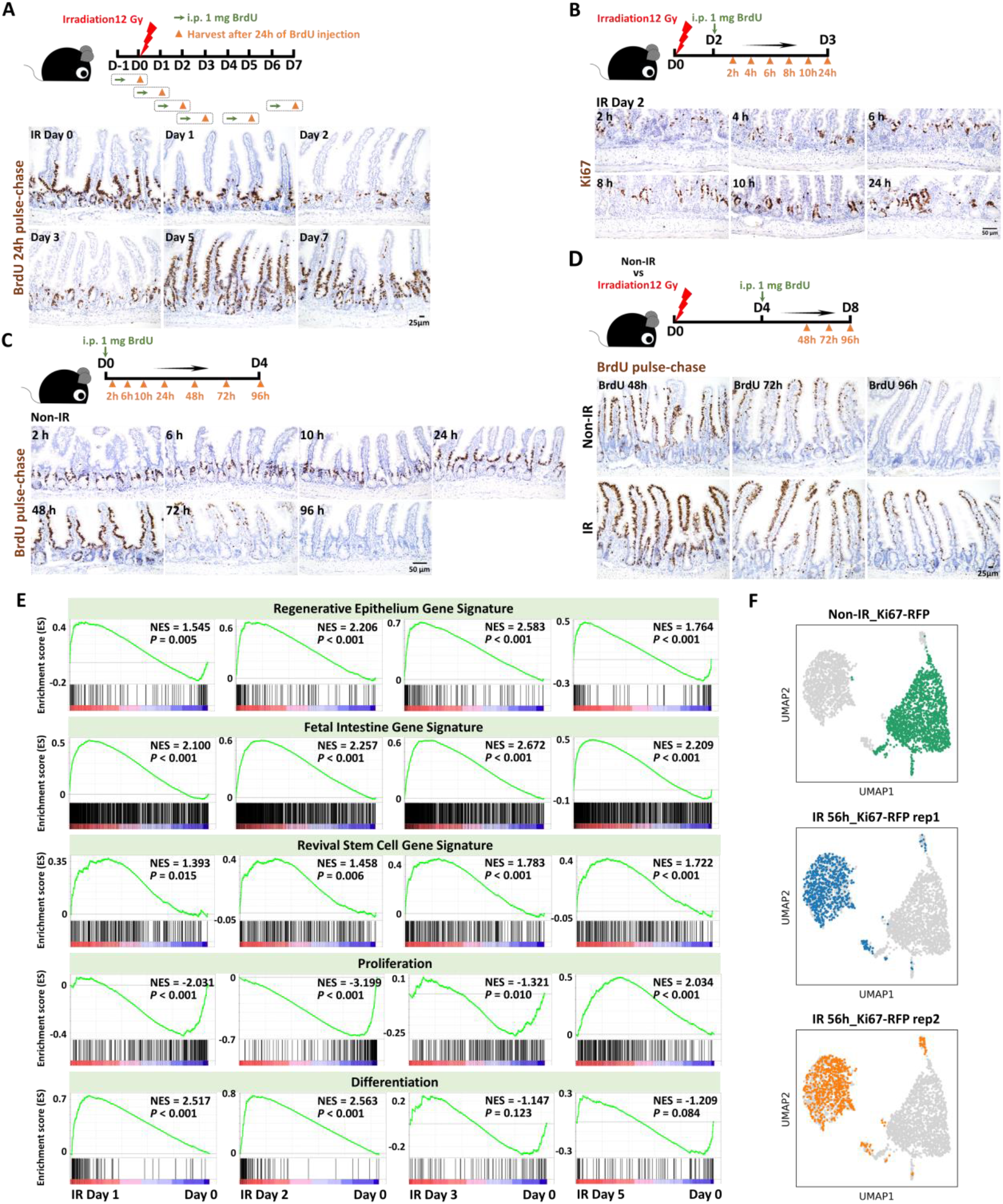
Day 2 to Day 3 post-irradiation is a critical regeneration period of crypt cells. **(A)** BrdU pulse-chase immunostaining of the intestine upon irradiation (representative of 3 biological replicates). Mice were injected with 1 mg of BrdU before or after irradiation. Intestines were collected after 24 hours of BrdU injection (see time-course details in the schematic of experimental design). **(B)** Immunostaining of Ki67 in the intestinal tissues of mice after 12 Gy irradiation. Ki67: proliferative marker, brown color; representative of 3 biological replicates. The schematic of experimental design was the same as BrdU experiment performed in **Figure 1B. (C)** BrdU pulse-chase immunostaining of the intestinal tissues collected from the non-IR control mice (representative of 3 biological replicates). For BrdU immunohistochemistry, mice were injected with 1 mg of BrdU. Intestinal tissues were harvested after 2, 6, 10, 24, 48, 72 and 96 hours of BrdU injection. **(D)** BrdU pulse-chase immunostaining of the intestinal tissues (representative of 3 biological replicates). Mice were injected with 1 mg of BrdU at Day 4 post-irradiation. Tissues were collected after 48, 72 and 96 hours of BrdU injection (see time-course details in the schematic of experimental design). BrdU positive cells are marked brown. Non-IR mice were used as controls. **(E)** GSEA examines expression changes in gene signatures of regenerative epithelium, fetal spheroid, revival stem cell, proliferation, and differentiation (Ayyaz et al., 2019; Merlos-Suarez et al., 2011; Mustata et al., 2013; Wang et al., 2019; Yui et al., 2018) at time points post-irradiation. Bulk RNA-seq was used in this analysis (GSE165157 (Qu et al., 2021), crypt cells, n=2 biological replicates per time-course, Kolmogorov-Smirnov test). **(F)** scRNA-seq reveals that sorted Ki67-RFP positive cells show different transcriptome profiles after 56 hours of irradiation compared to the non-IR controls. Number of cells in each condition was Non-IR Ki67-RFP positive cells: n=1739; IR 56h Ki67-RFP positive cells replicate 1: n=677; IR 56h Ki67-RFP positive cells replicate 2: n= 669. Ki67-RFP positive cells were isolated and sorted from crypt cells of *Mki67tm1*.*1Cle/J* mice after 56 hours of IR vs. non-IR control.

**Figure S2.**
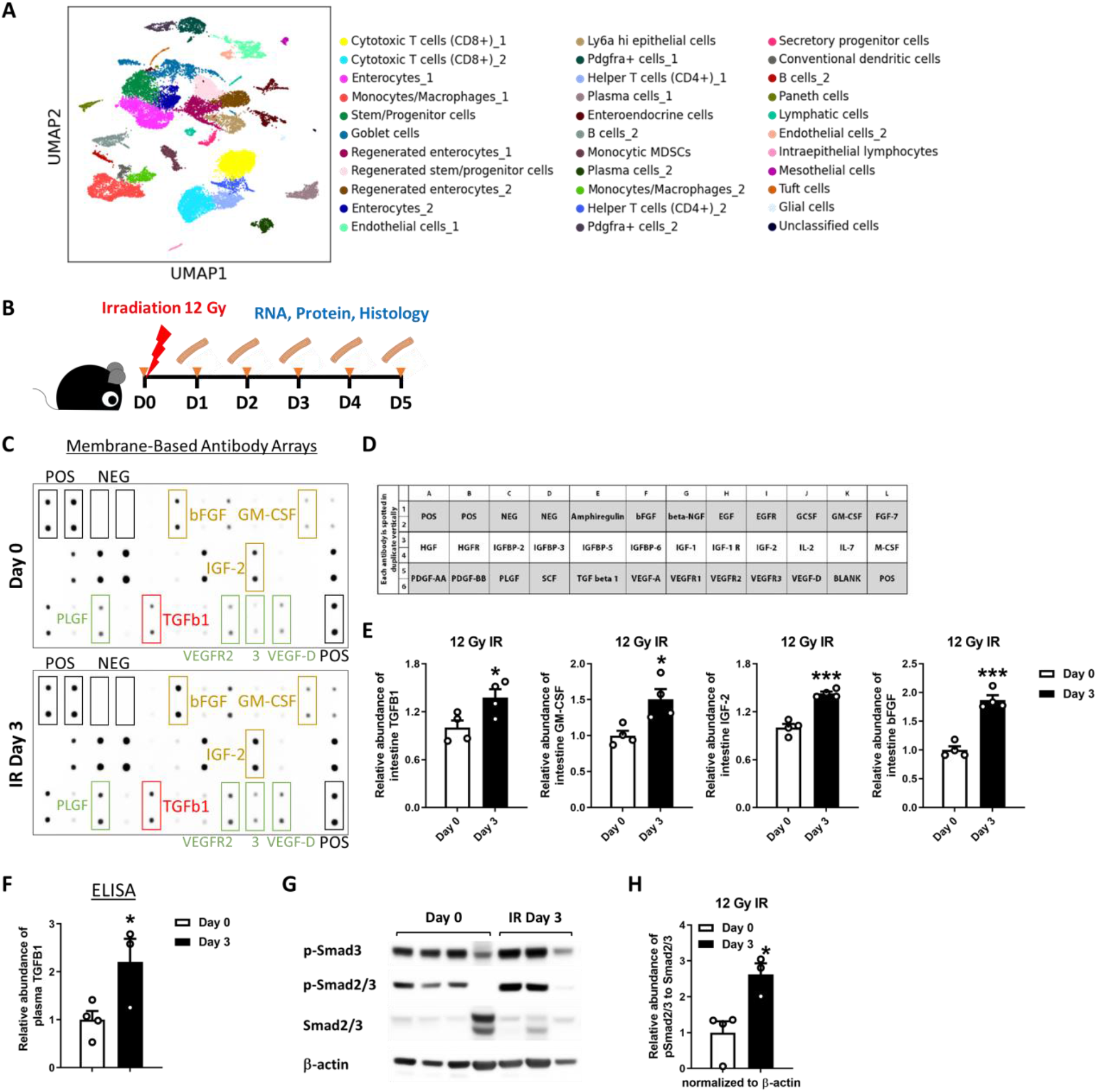
TGFB signaling is activated in Day 3 irradiated mouse intestine. **(A)** Identified cell populations of intestinal scRNA-seq data following mouse irradiation at days 0, 1, 3, 7, and 14, as shown in **Figure 2B** (GSE165318, n=3-4 biological replicates per time-course). **(B)** Schematics of IR time-course and sample collection for RNA, protein and histology. **(C-E)** IGF-2 is reported to preprogram maturing macrophages (TGFB1 producers) to acquire oxidative phosphorylation-dependent anti-inflammatory properties (Du et al., 2019). On the other hand, TGFB1 is known to enhance CM-CSF (Celada and Maki, 1992) and bFGF (Pertovaara et al., 1993), and we found TGFB1 and its related growth factors are all enriched in Day 3 intestine post-irradiation. Increased PLGF and VEGF were reported under pathologic situations, which are also observed in our study. **(E)** Quantification (n=2 independent experiments, 2 technical replicates per membrane, Student’s t-test at *P* < 0.001*** and *P* < 0.05*). **(F)** Increased TGFB1 is observed in plasma at Day 3 post-irradiation compared to Day 0, as determined by ELISA (n=3-4 biological replicates, Student’s t-test at *P* < 0.05*). **(G-H)** Western blot reveals that increased p-Smad3 and p-Smad2/3 levels (TGFB pathway) are detected in the intestine after 3 days of irradiation. **(H)** Quantification of western blot (n=3-4 biological replicates, Student’s t-test at *P* < 0.05*). TGFB1: Transforming growth factor beta 1; bFGF: Basic fibroblast growth factor; GM-CSF: Granulocyte-macrophage colony-stimulating factor; IGF-2: insulin-like growth factor; PLGF: Placental growth factor; VEGF: Vascular endothelial growth factor.

**Figure S3.**
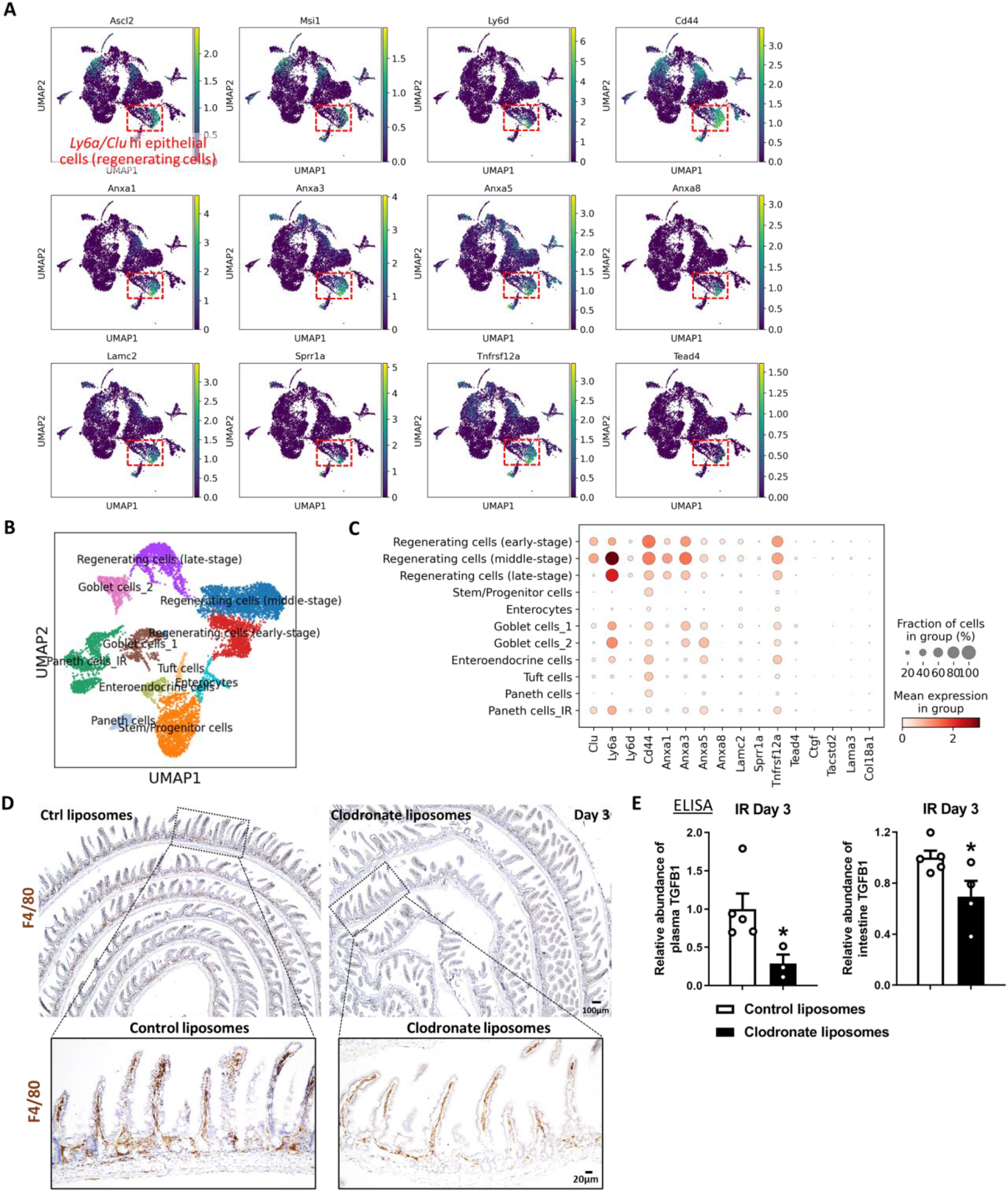
Fetal/regenerative genes are highly enriched during regeneration. **(A)** scRNA-seq of mouse intestines across a time-course post-irradiation (GSE165318). Of all the epithelial cells in the dataset (marked by *Epcam* expression in **Figure 3B**), regenerating cells expressing *Tgfbr2* are also highly enriched in transcript levels of reserve stem cell genes and fetal/regenerative marker genes. **(B-C)** *Msi1-CreERT2; R26-mTmG* mice were treated with tamoxifen for 15 hours, and *Msi1* positive GFP cells were sorted for scRNA-seq. Identified cell populations **(B)** of intestinal scRNA-seq data following mouse irradiation at days 0, 1, 2, 3, and 5 (GSE145866 (Sheng et al., 2020)). Fetal/regenerative transcripts **(C)** are highly enriched in these regenerating cells. **(D-E)** Monocytes/Macrophages were depleted using clodronate-containing liposomes (see schematic of experimental design in **Figure 3F**). Immunostaining **(D)** was performed to confirm reduction in F4/80 (monocyte/macrophage marker) expressing cells upon treatment. Reduced levels of TGFB1 are observed in the plasma and intestine upon treatment of clodronate liposomes compared to treatment with control liposomes, as evidenced by ELISA **(E)**. Plasma and duodenal fragments were collected 3 days post-irradiation (n=3-5 biological replicates, Student’s t-test at *P* < 0.05*).

**Figure S4.**
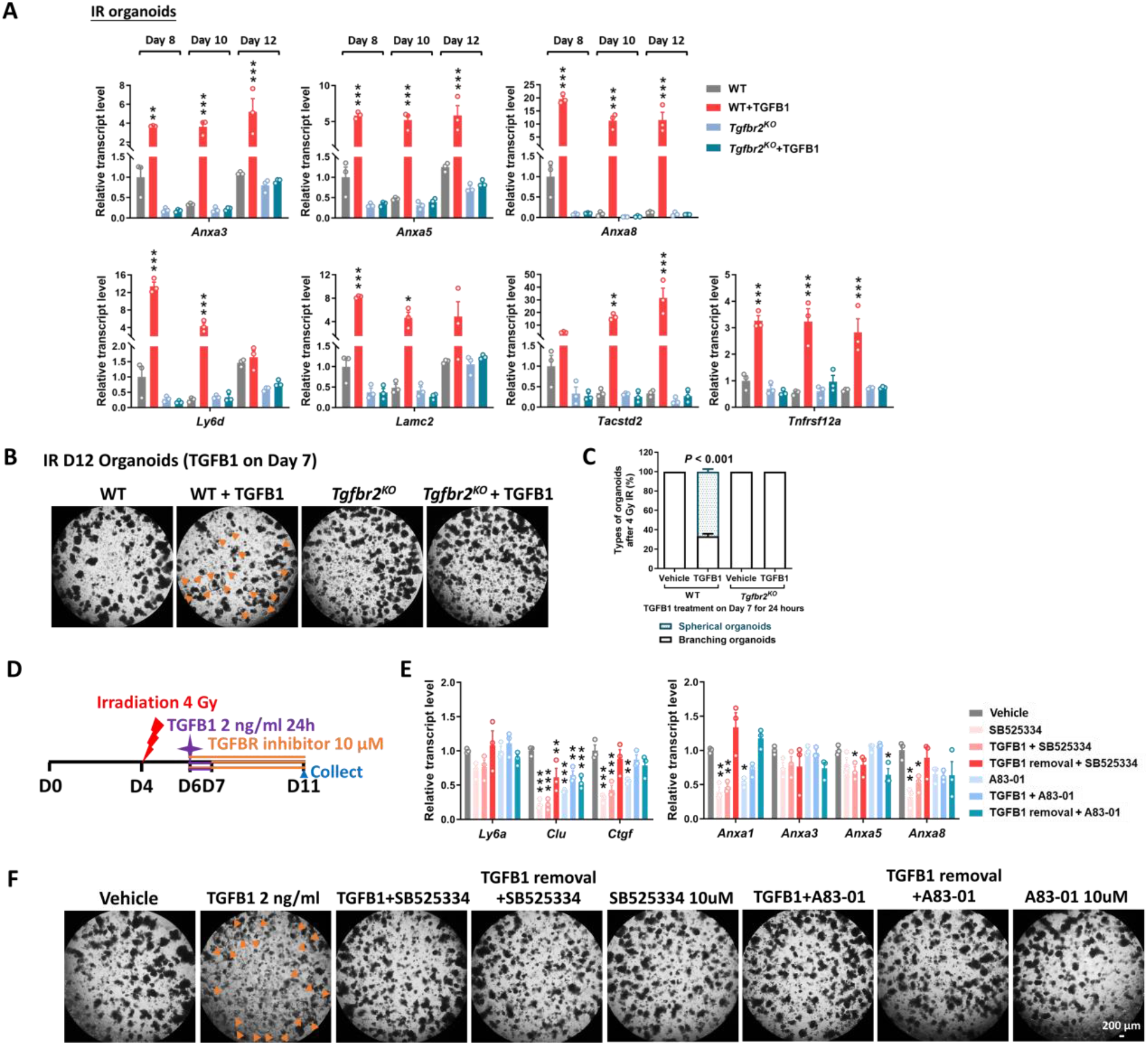
Inactivation of *Tgfbr2* abolishes TGFB1-induced spheroid morphology and fetal/regenerative gene signatures. **(A)** qRT-PCR indicates that expression of regeneration marker genes increases within 24 hours and remains elevated for at least 5 days post-TGFB1 treatment in organoids post-irradiation. This induction is dependent on *Tgfbr2*. Schematic of experimental design of qRT-PCR for panel A is depicted in **Figure 4F**. Transcript levels are relative to WT, and statistical comparisons were performed using one-way ANOVA followed by Dunnett’s post at *P* < 0.001***, *P* < 0.01** or *P* < 0.05* (n=3 independent organoid cultures). **(B-C)** Loss of *Tgfbr2* abolishes TGFB1 induced spheroid morphology (orange arrows, n=3 independent organoid cultures). **(B)** Representative images. **(C)** Quantification. Statistical comparisons were performed using one-way ANOVA followed by Dunnett’s post at *P* < 0.001. Schematic of experimental design for panels B-C is in **Figure 4F**. These findings were corroborated by adding TGFB receptor inhibitors **(D-F). (D)** Schematic of intervention experiment using TGFB receptor inhibitors. Primary intestinal organoids were exposed to 4 Gy of irradiation on Day 4, followed by TGFB1 treatment (2 ng/ml) on Day 6 for 24 hours. TGFB receptor inhibitor, 10 μM SB525334 or A83-01, was added at the same time with TGFB1 treatment, or added after removal of TGFB1, or added without TGFB1 pre-treatment. The corresponding intervention continued until Day 11. Organoids were imaged on Day 11, and then collected for qRT-PCR. **(E)** Presence of TGFB receptor inhibitors suppresses TGFB1-induced expression of fetal/regenerative genes. Transcript levels relative to vehicle, and statistical comparisons were performed using one-way ANOVA followed by Dunnett’s post at *P* < 0.001***, *P* < 0.01** or *P* < 0.05* (n=3 independent organoid cultures). All the qRT-PCR data are presented as mean ± SEM. **(F)** TGFB receptor inhibitors abolish TGFB1-induced spheroid morphology (orange arrows, n=3 independent organoid cultures).

**Figure S5.**
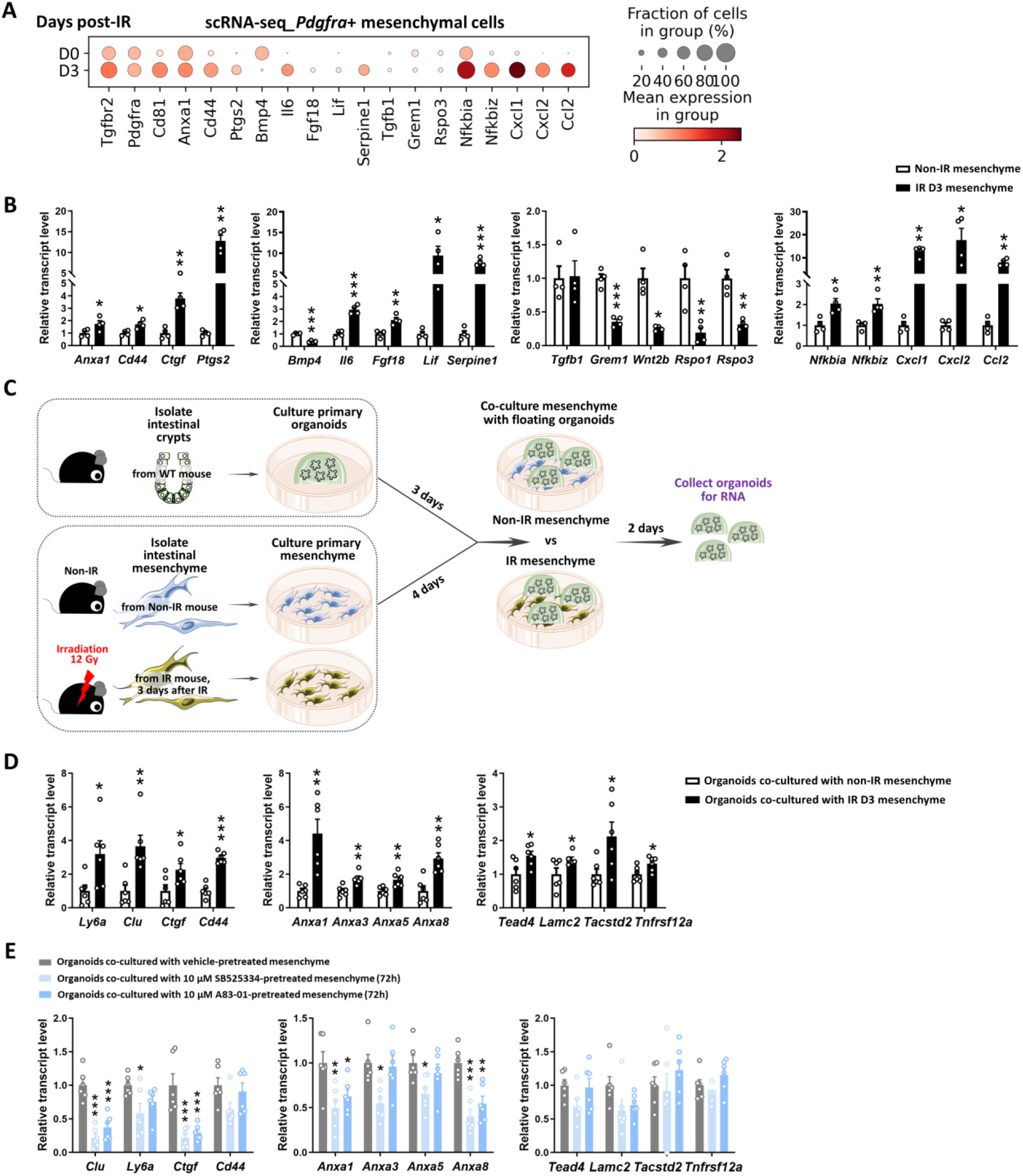
Mesenchyme isolated from mice post-irradiation promotes fetal-like gene signatures of intestinal organoids. **(A-B)** Cultured mesenchymal cells isolated from mice with or without irradiation harbor similar features of cells isolated from primary *Pdgfra* positive mesenchymal cells, as evidenced by scRNA-seq dot plots **(A)** of *Pdgfra* positive mesenchyme cell cluster (primary cells, GSE165318) and qRT-PCR **(B)** of cultured mesenchyme (n=4 independent mesenchyme cultures with 2 different primary cell lines). **(C)** Schematic of co-culture experimental design. Intestinal mesenchymal cells were isolated from mice with or without irradiation (12 Gy, 3 days post-IR) and cultured for 4 days. They were then overlaid with Day 3 primary organoids in matrigel bubbles for another 2 days. Co-cultured organoids were collected as floating matrix bubbles for qRT-PCR. **(D)** qRT-PCR reveals that compared to the non-IR condition, mesenchyme isolated from mice post-irradiation shows stronger induction of fetal/regenerative gene expression in co-cultured organoids (n=6 independent organoid cultures with 2 different cell lines of mesenchyme and 3 different cell densities). The qRT-PCR data are presented as mean ± SEM (Student’s t-test at *P* < 0.001***, *P* < 0.01** or *P* < 0.05* for panels B and D). **(E)** Mesenchyme pre-treated with TGFBR inhibitors suppresses fetal-like gene signatures of co-cultured intestinal organoids. Passaged intestinal mesenchyme cells were pre-treated with vehicle or TGFBR inhibitors for 3 days, and then co-cultured with Day 3 primary organoids for 2 days (similar procedure as shown in **Figure 5D**). Co-cultured organoids were collected for qRT-PCR (n=6 independent organoid cultures with 2 different cell densities of mesenchyme). TGFBR inhibitors were removed during co-culture. Transcript levels relative to vehicle control, and statistical comparisons were performed using one-way ANOVA followed by Dunnett’s post at *P* < 0.001***, *P* < 0.01** or *P* < 0.05*.

**Figure S6.**
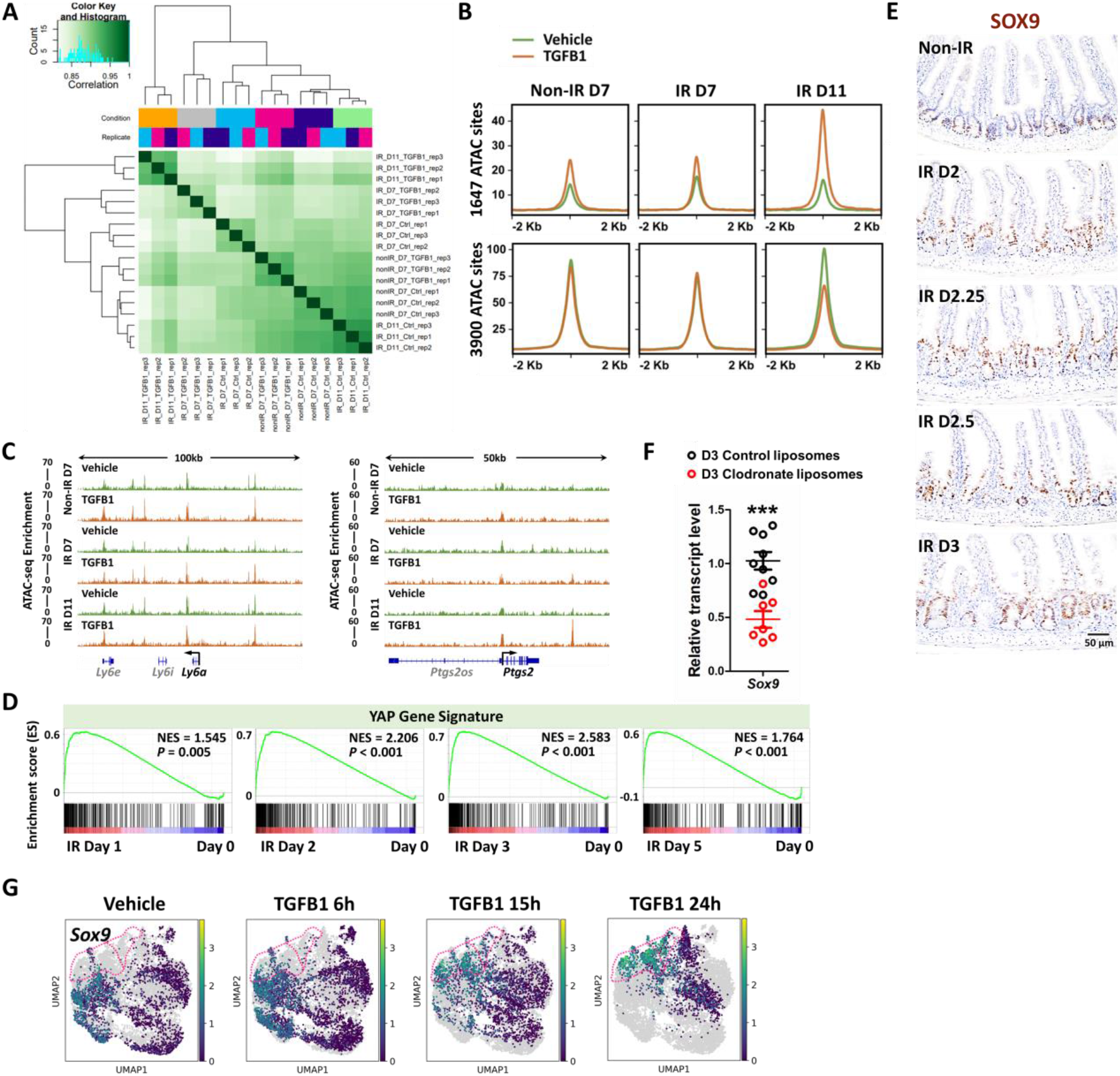
TGFB1 activates intestinal YAP and SOX9 regenerative programs. **(A)** ATAC-seq correlation heatmap of organoids upon TGFB1 treatment (DiffBind package, n=3 independent organoid cultures, at MACS2 called peaks of q < 0.05). The experimental design is the same as bulk RNA-seq and shown in **Figure 4A. (B)** Average signal of ATAC-seq in accessible chromatin regions of vehicle or TGFB1 treated organoids (Differential ATAC-seq regions defined in **Figure 6A**, Diffbind FDR < 0.01). **(C)** Examples of genes located at ATAC-seq enriched regions of TGFB1-treated organoids using IGV. **(D)** GSEA reveals gene signatures of elevated YAP signaling (Gregorieff et al., 2015) post-irradiation, using a bulk RNA-seq of an IR time-course data set (GSE165157 (Qu et al., 2021), crypt cells, n=2 biological replicates per time-course, Kolmogorov-Smirnov test). **(E)** Intestinal immunostaining of SOX9 from day 2 to day 3 post-IR vs. non-IR (brown color; representative of 3 biological replicates). **(F)** Depletion of monocytes/macrophages (main cell sources of TGFB1 secretion) results in a downregulation of *Sox9*, as evidenced by qRT-PCR (n=7-9 biological replicates, duodenal fragments, Student’s t-test at *P* < 0.001***). **(G)** scRNA-seq UMAP plots indicate that TGFB1 induces *Clu*-expressing cells that highly express *Sox9* (shown within pink dotted line, defined in **Figure 4K**). Number of cells in each condition was Vehicle: n= 2815; TGFB1 6h: n= 4071; TGFB1 15h: n= 2788; TGFB1 24h: n=2177.

**Figure S7.**
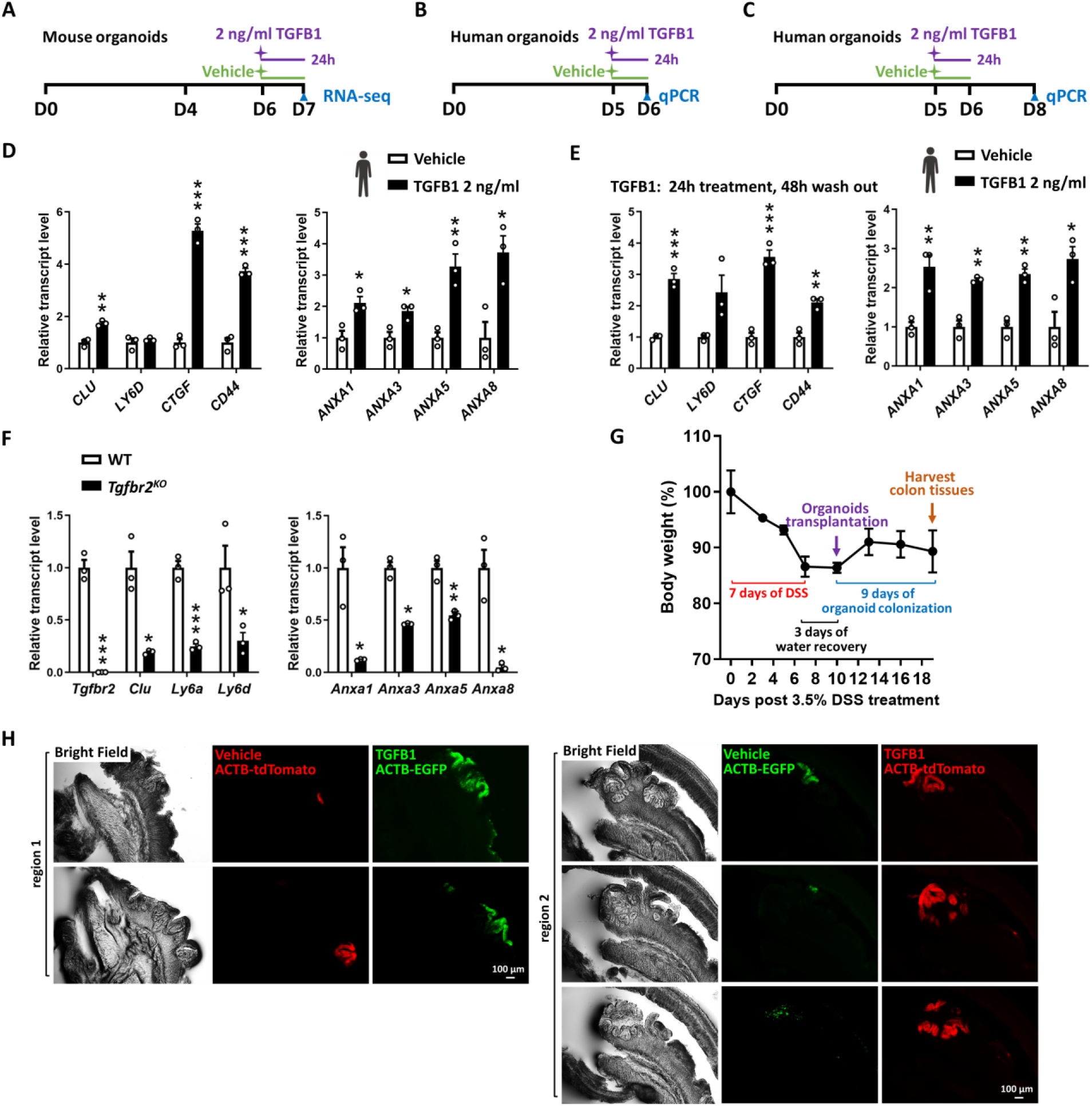
TGFB1 pre-treated organoids transplant more efficiently in DSS-damaged colon. **(A)** Schematic for the bulk RNA-seq experimental design of non-IR mouse organoids. **(B-E)** TGFB1-induced fetal/regenerative gene signatures are conserved in adult human duodenal organoid cultures. qRT-PCR indicates that expression of regeneration marker genes increases and remains elevated for at least 3 days post-TGFB1-treatment in adult human duodenal organoid cultures. TGFB1 treatment was performed for a 24-hour window either **(B, D)** 24 hours or **(C, E)** 72 hours before collection of cells. Human organoids were treated with TGFB1 on Day 5 (n=3 independent organoid cultures, Student’s t-test at *P* < 0.001***, *P* < 0.01** or *P* < 0.05*). All the qRT-PCR data are presented as mean ± SEM. **(F)** To inactivate*Tgfbr2, Tgfbr2f/f*; *Villin-Cre*^*ERT2*^ mice were injected with tamoxifen for 4 consecutive days, and harvested 4 days after the first injection for primary organoid culture. Littermate controls were injected with vehicle. WT and *Tgfbr2*^*KO*^ organoids were collected on Day 7. qRT-PCR reveals that fetal/regenerative genes are suppressed in intestinal organoids upon loss of *Tgfbr2* (n=3 independent organoid cultures, Student’s t-test at *P* < 0.001***, *P* < 0.01** or *P* < 0.05*). **(G)** Body weight of mice treated with DSS in organoid transplantation experiment. NOD mice were treated with 3.5% DSS for 7 days. After 3 days of water recovery, organoid transplantation was performed and colon tissues were harvested 9 days after organoid transplant (n=5 biological replicates). **(H)** TGFB1 pre-treatment enhances organoid engraftment in DSS-treated mice. Examples of representative regions. Consecutive cryosections that represent the same colonization event were counted as one region. 21 organoid-colonized regions were found from 5 mice.

